# LARP6 regulates the mRNA translation of fibrogenic genes in liver fibrosis

**DOI:** 10.1101/2025.01.16.633226

**Authors:** Hyun Young Kim, Orel Mizrahi, Wonseok Lee, Sara B. Rosenthal, Cuijuan Han, Brian A. Yee, Steven M. Blue, Jesiel Diaz, Jyotiprakash Jonnalagadda, Kanani Hokutan, Haeum Jang, Chen-Ting Ma, Andrey Bobkov, Eduard Sergienko, Michael R. Jackson, Branko Stefanovic, Tatiana Kisseleva, Gene W. Yeo, David A. Brenner

## Abstract

Metabolic syndrome and excessive alcohol consumption result in liver injury and fibrosis, which is characterized by increased collagen production by activated Hepatic Stellate Cells (HSCs). LARP6, an RNA-binding protein, was shown to facilitate collagen production. However, LARP6 expression and functionality as a regulator of fibrosis development in a disease relevant model remains elusive. By using snRNA-sequencing, we show that LARP6 is upregulated mainly in HSCs of liver fibrosis patients. Moreover, LARP6 knockdown in human HSCs suppresses fibrogenic gene expression. By integrating eCLIP analysis and ribosome profiling in HSCs, we show that LARP6 interacts with mature mRNAs comprising over 300 genes, including RNA structural elements within *COL1A1*, *COL1A2*, and *COL3A1* to regulate mRNA expression and translation. Furthermore, LARP6 knockdown in HSC attenuates fibrosis development in human liver spheroids. Altogether, our results suggest that targeting LARP6 in human HSCs may provide new strategies for anti-fibrotic therapy.

**Highlights:**

- LARP6 is upregulated in liver fibrosis, mainly in HSCs.
- LARP6 knockdown in human HSCs reduces liver fibrosis development.
- Of the hundreds of gene targets, LARP6 interacts most with collagen mRNAs.
- LARP6 regulates mRNA translation via interaction with 5’UTRs.

## Introduction

Chronic liver injury from various etiologies leads to the development of liver fibrosis, and the stage of liver fibrosis correlates closely with patient mortality. The most prevalent chronic liver disease in western countries is metabolic dysfunction-associated steatotic liver disease (MASLD)^1^. MASLD ranges from simple steatosis to metabolic dysfunction-associated steatohepatitis (MASH) which features lobular inflammation, Hepatic Stellate Cell (HSC) activation, and fibrosis. Liver fibrosis is characterized by excessive deposition of extracellular matrix (ECM) proteins, especially Collagen Type I and III, produced by activated HSCs^2,3^. Therefore, targeting HSC activation or collagen production is the primary strategy to halt fibrosis and improve clinical outcomes.

Collagen Type I and III have an evolutionary conserved 5’ stem loop (5’SL) mRNA structure^4^. Interestingly, mice with a knock-in mutation introduced into the 5′SL of collagen α1 demonstrated a decrease in liver fibrosis development and a lower expression of collagen in HSCs and mouse embryonic fibroblasts, indicating the importance of the 5’SL structure to collagen expression in HSCs and fibrosis^5^.

Previous screens for regulators of collagen expression revealed that the RNA-binding protein (RBP) La-related protein 6 (LARP6) interacts *in vitro* with the collagen 5’SL in cytoplasmic fraction of human fibroblasts by electromobility shift assay. LARP6 was also found to bind non-collagen stem-loop RNA structures with multiple motifs, implying structural specificity rather than sequence specificity^6^. At the cellular level, reducing LARP6 expression results in a decrease of collagen production, indicating that LARP6 regulates collagen expression and contributes to liver fibrosis development^4,7^. However, while liver fibrosis is characterized with extensive changes at the transcriptomic and proteomic levels, a comprehensive analysis of LARP6 RNA targets in a disease relevant model remains unexplored^8–10^.

LARP6 is well conserved in evolution, with genetic elements that are conserved in almost all eukaryotes. It is part of the LA and Related Proteins (LARPs) superfamily, which is composed of the RNA binding domains La Motif (LaM) and RNA Recognition Motif (RRM)^11^. Interestingly, an additional motif is found at the LARP6 C-terminus that is suggested to mediate interaction with mRNA substrates^12,13^. However, the post-transcriptional effects of LARP6-mRNA interaction remains unclear. LARP6 was shown to interact with collagen mRNA to enhance their expression, presumably via increase in their mRNA stability and translation^4^. Interestingly, previous individual-nucleotide resolution UV crosslinking and immunoprecipitation (iCLIP) of exogenously expressed LARP6 in breast cancer cells revealed that LARP6 interacts with several thousands of genes. Those interactions reside mostly within protein coding genes and are enriched next to the mRNA translation initiation site. Moreover, LARP6 was suggested to interact with ribosomal mRNAs and mediate their sub-cellular localization and translation^14^. However, unbiased and accurate evaluation of direct LARP6 effect on mRNA expression and translation at the endogenous levels remained absent.

Here, we investigated the LARP6-dependent mechanism in liver fibrosis among patients with MASH and metabolic dysfunction-associated Alcohol-related liver disease (MetALD). We identified that LARP6 is highly expressed in activated HSCs from human fibrotic livers. We demonstrated that fibrogenic genes are decreased in LARP6 knockdown cells at the mRNA and protein levels. This effect was further enhanced in HSCs activated by TGFβ1, a key fibrogenic cytokine driving HSC activation during the progression of chronic liver disease.

By applying enhanced crosslinking and immunoprecipitation (eCLIP) analysis and ribosome profiling in human HSCs, we captured endogenous LARP6-specific binding and regulation of collagen mRNAs as the most enriched RNA-binding targets of LARP6. Beyond these targets, we show that LARP6 binds mRNAs from hundreds of different genes. We also demonstrated a list of newly identified targets in collagen-related pathways that are regulated by LARP6 both at the mRNA translation and expression levels, thus expanding the knowledge of its binding repertoire and regulatory functions. Lastly, we demonstrated in 3D human liver spheroids that HSC-specific LARP6 knockdown suppresses fibrosis development by reducing collagen production. Altogether, the RNA regulatory network influenced by LARP6 suggests that targeting LARP6 in HSCs holds promise for anti-fibrotic therapy.

## Results

### LARP6 is upregulated in activated HSCs from MASH and MetALD human livers

We compared the gene expression and chromatin accessibility profiles of HSCs from human NORMAL (n=5), MASL (n=4), MASH (n=3), and MetALD (n=6) livers at single-cell resolution^15^ (Figure **1A**). Deidentified human donor livers were examined by a pathologist, and livers with a MASH clinical research network (MASH/CRN) score of 0 were diagnosed as NORMAL (donors D1-5). Livers with steatosis and without fibrosis were identified as MASL (donors D6-9), while those with steatosis, inflammation, and fibrosis were diagnosed as MASH or MetALD (donors D10-18). Based on patient history, human livers without an alcohol consumption history were identified as MASH (donors D10-12), and those with a significant history of alcohol consumption (>2 drinks per day) were identified as MetALD (donors D13-18, Figure **1A**, Figure **S1A**). No significant difference in liver fibrosis was observed between the MASH and MetALD groups, as shown by fibrosis stage and the Sirius Red staining positive area (Figures **1B-C**, Figure **S1B**). We performed single nucleus (sn)RNA-sequencing and snATAC-sequencing of isolated nuclei from snap-frozen liver tissues of these donors.

**Figure 1.**
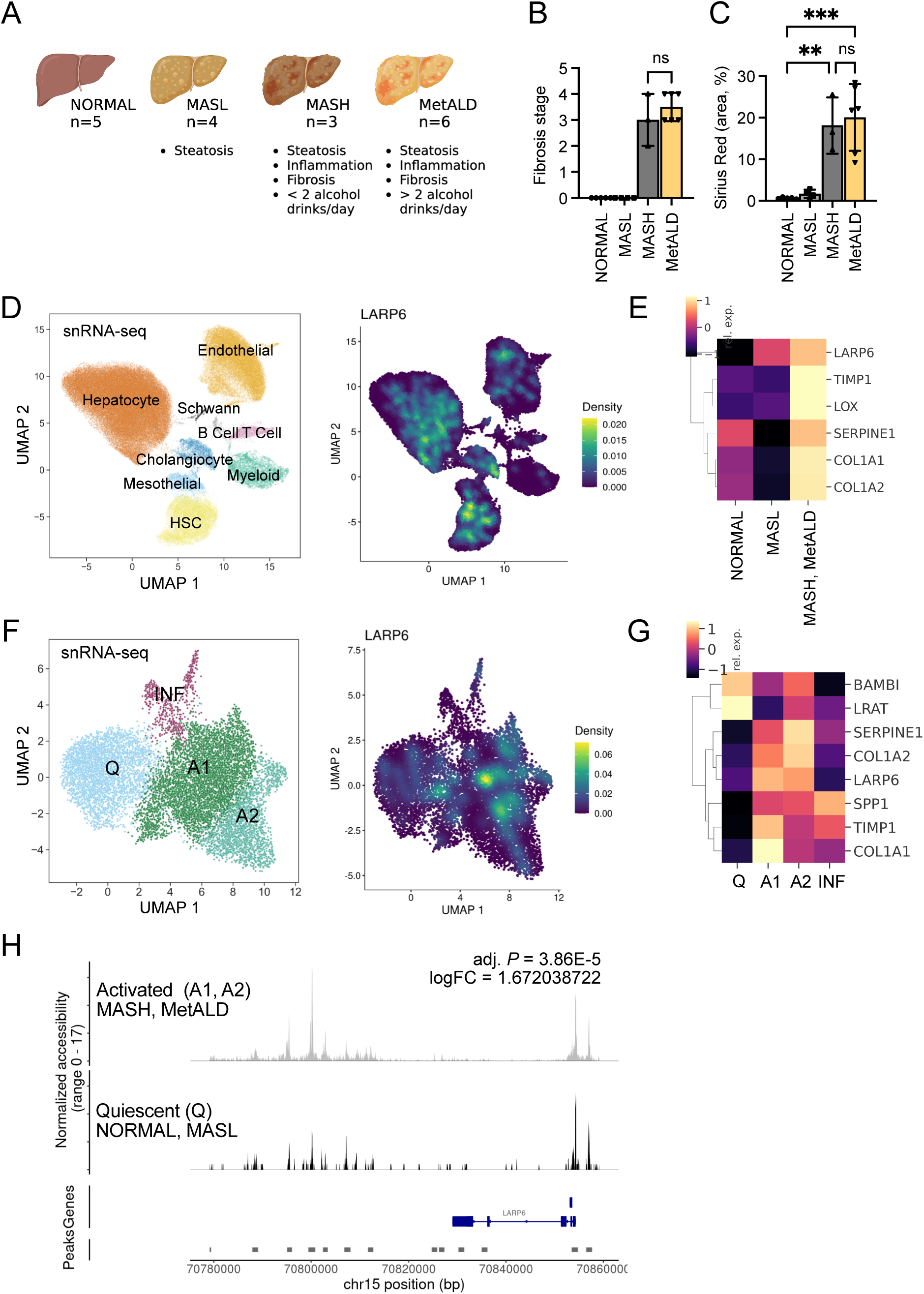
LARP6 is upregulated in activated HSCs from human fibrotic livers. **(A)** Schemetic illustration of the human livers selected for snRNA-seq and snATAC-seq. Four distinct liver diagnoses of NORMAL, MASL, MASH, and MetALD were defined by a combination of pathological examination and alcohol consumption history. **(B)** Fibrosis scores were graded by a pathologist in a double-blinded manner. **(C)** Livers were stained for Sirius Red, and the positive area was calculated as a percentage. Data are presented as mean ± SD; \*\**P* < 0.01, and \*\*\**P* < 0.001; ns, not significant; one-way ANOVA followed by Tukey’s test. **(D)** SnRNA-seq UMAP plot of the integrated dataset of liver cells from all donors, showing identified cell types (left) and color-coded UMAP for LARP6 expression (right). **(E)** Heatmap representing relative expression of LARP6 and selected genes specific for MASH and MetALD HSCs. **(F)** 1F: HSCs were clustered according to gene expression profile using Seurat v. 4.0. SnRNA-seq UMAP plot of the integrated dataset of HSCs from all donors, showing four HSC clusters (left); quiescent (Q), two activated (A1 and A2), and inflammatory (INF), and the UMAP color-coded for LARP6 smoothed expression (right), using Nebulosa for expression smoothing^51^. **(G)** Heatmap representing relative expression of LARP6 and fibrogenic genes across HSC clusters. **(H)** snATAC-seq normalized accessibility peaks of LARP6. Chromatin accessibility between the peak upstream of LARP6 (peak = chr15- 70799442-70800783) was calculated in activated HSCs from fibrotic livers (MASH, MetALD) and quiescent HSCs from nonfibrotic livers (NORMAL, MASL).

NORMAL, MASL, MASH, and MetALD liver datasets were integrated, clustered, and different cell types were identified by marker genes (*NGFR*^3^, *CYGB*^16^, *COL1A1*, *RBP1*^17^, and *HGF*^18^) and PanglaoDB database^15,19^. Based on our snRNA-seq analysis, *LARP6* was mainly expressed in HSCs, the major Collagen Type I-producing myofibroblasts in fibrotic liver (Figure **1D**, Figure **S1C**). Especially, *LARP6* was upregulated in HSCs from MASH and MetALD livers, along with fibrogenic genes including *COL1A1*, *COL1A2,* and *TIMP1* (Figure **1E**). We subclustered HSCs using a previously identified set of marker genes^20^ to study the characteristics of *LARP6*-positive HSCs. In the snRNA-seq dataset, we found four subclusters of HSCs; quiescent (Q), activated 1 (A1), activated 2 (A2), and inflammatory (INF) HSCs. *LARP6* was highly expressed in A1 and A2 HSCs (Figures **1F-G**, Figure **S1D**). Chromatin accessibility near or within the loci of *LARP6* gene increased in activated HSCs from MASH or MetALD livers (logFC=1.67, adj.*P*<4E-5, Figure **1H**). A database for transcription factor-target gene relationship^21^ showed that *LARP6* was co-regulated by a group of ECM-specific gene-regulating transcription factors (red triangles), including *FOXP1* and *FOXA1*^22^ (Figure **S1E)**.

### Knockdown of LARP6 decreases TGFβ1-responsive genes in HSCs

LARP6 protein expression was assessed in human liver tissues. Human liver tissues were stained with anti-LARP6 antibody and counterstained with hematoxylin (Figure **2A**). LARP6 expression significantly increased in MASH and MetALD human livers (Figure **2B**). In human MetALD livers, cells that stained positive for LARP6 were also positive for the myofibroblast marker, α-smooth muscle actin (αSMA), in serial sections (Figure **2C**). This indicates that most of the LARP6-positive cells are activated HSCs.

**Figure 2.**
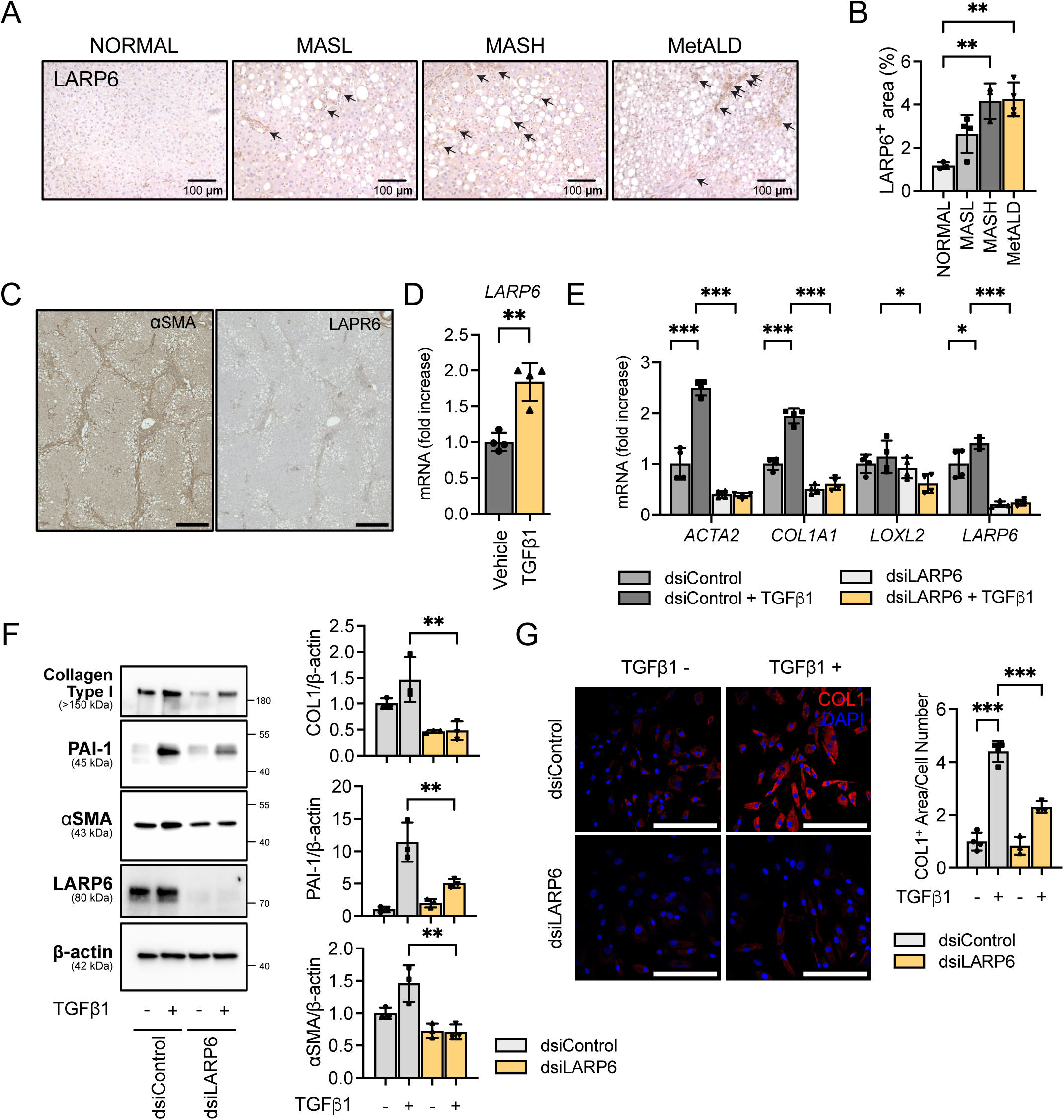
Knockdown of LARP6 inhibits HSC activation. **(A)** Human liver tissues were stained with an anti-LARP6 antibody. **(B)** The LARP6-positive area was calculated as a percentage. **(C)** LARP6-positive cells stained positive for HSC marker αSMA on serial sections of human fibrotic liver (donor D13). **(D)** Cultured human HSCs (donor D19) were stimulated with human TGFβ1 (5 ng/ml) for 24 h. **(E-G)** HSCs (donor D19) were transfected for 48 h with dsiRNA and stimulated with human TGFβ1 (5 ng/ml) for 24 h. The expression of fibrogenic genes was measured in LARP6-targeting dsiRNA (vs dsi-negative control)-transfected human HSCs ± TGFβ1 at the **(E)** mRNA and **(F)** protein levels. **(G)** dsiRNA-transfected HSCs ± TGFβ1 were stained with an anti-Collagen Type I(COL1A1) antibody, and the COL1-positive area was calculated and normalized by the cell number counted using DAPI. Data are presented as mean ± SD (n=3 or 4); \**p* < 0.05, \*\**p* < 0.01, and \*\*\**p* < 0.001, one-way ANOVA followed by Tukey’s test.

Functional properties of LARP6 were evaluated in human HSCs isolated from MASH or MetALD donors (donor D19-D21, Figure **S2A**). LARP6 expression was induced in response to treatment with human TGFβ1, the most potent activator of HSCs (Figure **2D**). Then, the role of LARP6 gene in HSC activation was assessed using LARP6-targeting dsiRNA (Dicer-Substrate Short Interfering RNAs; dsiLARP6). HSCs transfected with dsiLARP6 showed >90% gene knockdown efficiency compared to HSCs transfected with dsiRNA negative control (Figure **S2B**). Knockdown of LARP6 significantly downregulated fibrogenic genes *ACTA2* and *COL1A1* in TGFβ1-stimulated human HSCs (Figure **2E**, Figure **S2C, F**). LARP6 knockdown resulted in a significant reduction in fibrogenic markers, especially Collagen Type I expression (Figure **2F****, G** and Figure **S2D-E, S2G**), consistent with the role of LARP6 in regulating fibrotic collagens.

### LARP6 binds structural elements in TIS of collagen-associated genes

To comprehensively discover the direct RNA binding targets of LARP6 in human HSCs, we performed two biologically independent replicate eCLIP^23^ experiments on HSCs stimulated with or without TGFβ1 (Figure **3A** and Figure **S3A**). Using the CLIPper analysis workflow^24^, we captured 1,008 binding sites (peaks) enriched 8-fold over input (*P* < 0.001) within 397 genes. These binding sites mostly reside within 5’UTR, CDS and 3’UTR of mature mRNAs, comprising 7.66%, 48.06% and 31.54% of peaks in TGFβ1-stimulated HSCs, respectively (Figure **3B** and Table **S1**). A similar distribution was observed with eCLIP peaks from the unstimulated HSCs (Figure **S3B**) but with ∼38% reduction in total number of peaks (627), indicating an increased number of LARP6 binding sites with TGFβ1 treatment (Table **S1**), despite a comparable number of total sequenced reads. Gene ontology (GO) analysis on the genes containing significantly enriched peaks observed revealed statistically significantly (FDR<0.05) enriched terms associated with collagen-related pathways including collagen fibril organization, collagen metabolic process, ECM assembly and cell-matrix adhesion (Figure **3C** and Figure **S3C**).

**Figure 3.**
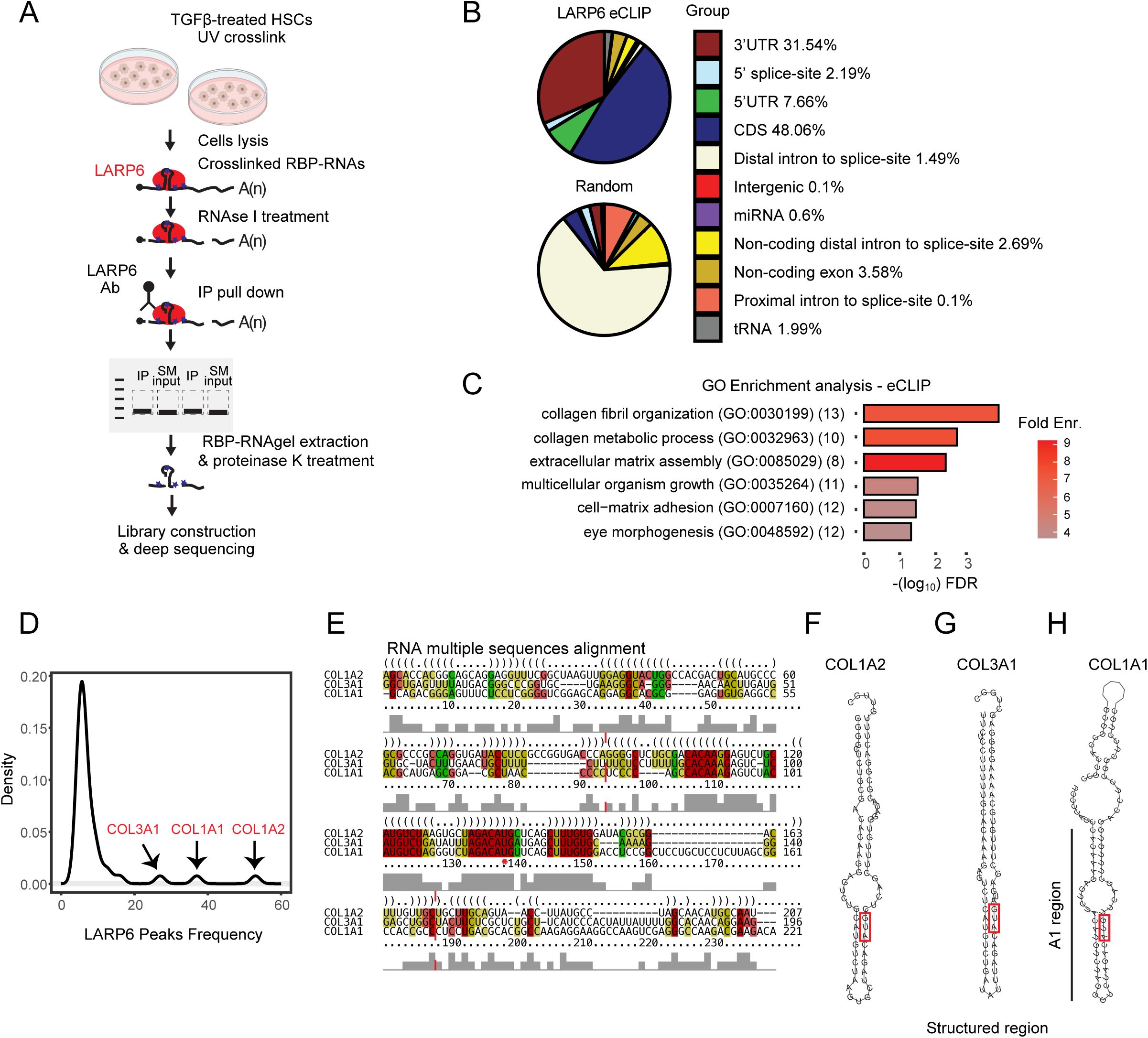
LARP6 interacts with mature collagen mRNAs. **(A)** Schematic illustration of eCLIP on LARP6 in TGFβ1-stimulated human HSCs. **(B)** LARP6 eCLIP binding analysis in TGFβ1-stimulated HSCs (Top pie chart; donor D22), and background distribution (low pie chart) of peaks. The percentages at the group-colored index represent binding distribution of the LARP6 eCLIP binding. **(C)** GO enrichment analysis using significant eCLIP peaks. RNA-seq genes with a minimum of 30 reads were used as background for enrichment analysis. **(D)** Density of eCLIP analysis of LARP6 targets with a minimum of five peaks. Peaks in *COL3A1*, *COL1A1*, and *COL1A2* are indicated with black arrows. **(E)** RNA multiple sequence alignment of the 5’UTR and the first coding exon for *COL1A2*, *COL3A1*, and *COL1A1* were calculated using LocARNA. Solid color represents conserved base-pairing with similar nucleotides (red), with two (yellow), or three (green) identities. The red asterisk represents the AUG of the canonical translation initiation site. Dashed lines flank the structured region that overlaps with the eCLIP signal. Grey columns represent sequence similarity per position in the multiple sequence alignment. **(F-H)** Structured region prediction using RNAfold for **(F)** *COL1A2*, **(G)** *COL3A1*, and **(H)** COL1A1. AUG of canonical translation initiation sites are indicated with red rectangles.

Next, we examined the frequency of LARP6 binding at the mRNA level. While almost half (47.6%) of the gene targets contained one binding site, the three genes that harbored the most sites are *COL1A2* (53 peaks), *COL1A1* (37) and *COL3A1* (27) in TGFβ1-stimulated HSCs, in consistent with previous *in vitro* LARP6 binding measurements^25^. Nevertheless, we observed that LARP6 binding does extend beyond these targets, with a total of 41 gene targets with at least five peaks in mRNAs in TGFβ1-stimulated HSCs and 20 targets in unstimulated cells (Figures **3D** and Figure **S3D**). The conserved regions in *COL1A1*, *COL1A2* and *COL3A1* that span their 5’UTR and first coding exon have the highest enrichment (reads from immunoprecipitation over size-matched input) among all the detected binding sites (log2 fold-change over input of ∼7.6, ∼7.87, and 9.3, respectively) together with newly identified interaction sites in CCNI 5’UTR (8.9) and LRP1 CDS (7.83). As previously shown, this region forms a stem-loop structure bound by LARP6 at the bulge regions. Interestingly, the eCLIP read coverage shows dips at the bulge and relatively low coverage at the hairpin region (Figure **S4**). As the binding seems to occupy a region wider than previously annotated, we examined the extent of sequence conservation of the collagen targets within the 5’UTR and the first exon. These regions share some nucleotide sequences, but its RNA structure is preferentially preserved at these coordinates (Figures **3E-H**). Strikingly, the most conserved sequences flank the start codon (AUG) of the translation initiation site (TIS), suggesting a common mechanism by which the structures engage with translational machinery (Figure **S5**). The structure conservation index (SCI) estimates structure similarity of different sequences to a consensus structure. Indeed, the most conserved subsection (Figure **3E**, marked with dashed red lines) consists of a higher SCI score (SCI = 0.7535), indicating a more conserved structure relative to the entire 5’UTR and first exon region (SCI = 0.4403).

Next we determined the evolutionary conservation scores (phyloP100way) in the flanking 30 bp window on either side of the start codon for the expressed genes in our data. The collagen targets (i.e. *COL1A1*, *COL1A2* and *COL3A1)* consist of a significantly well conserved region relative to random genes in the same relative coordinates flanking the canonical TIS (collagen targets conservation score ≥0.87, Figure **S3E**), indicating a well conserved regulatory element at the TIS of these collagen genes that mediated via a direct interaction with LARP6.

### TR-FRET assay and ITC validate the binding of LARP6

To validate our new LARP6 targets and to determine the strength of interaction between LARP6 and its RNA substrates independent of the cellular machinery, we designed RNA oligos from: 1) a region that flanks the canonical AUG translation initiation site (TIS; *COL1A1*-TIS) 2) 5’UTR region (i.e *CCNI*), and 3) spliced regions (i.e *COL5A1* and *LRP1*; Figure **4A**). We then measured the binding affinity of LARP6 with the target RNAs. We performed a TR-FRET assay using DIG-labeled 5’ SL structure flanking canonical AUG TIS of *COL1A1* (A1 RNA), which was shown to bind with LARP6 in cells^4^ (Figure **3H**). *IFIT1*-5’UTR RNA oligo which has not shown in eCLIP but fairly expressed in RNA-sequencing data, was used as a negative control.

**Figure 4.**
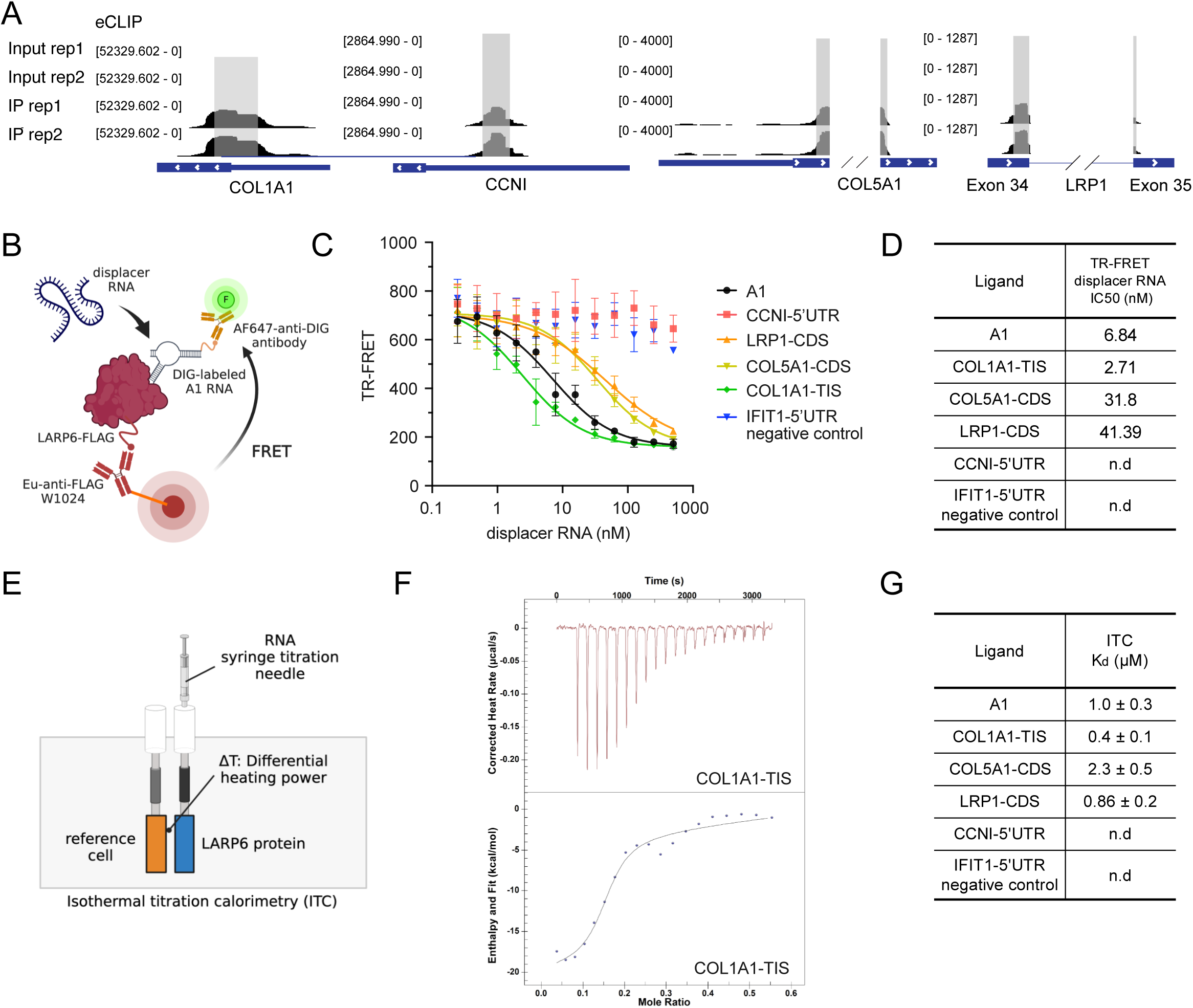
TR-FRET assay and ITC confirm the binding of LARP6. **(A)** LARP6-binding RNA regions (60 nucleotides) identified by eCLIP were synthesized for TR-FRET and ITC assays. **(B)** Schematic illustration of TR-FRET-based LARP6 binding assay. **(C)** Competition of LARP6 binding RNAs to DIG-labeled A1 RNA. Data are expressed as mean ± SD of duplicate. **(D)** Half-maximal inhibitory concentration (IC50) values of LARP6 binding to target RNAs; n.d, not detected. **(E)** Schematic illustration of ITC-based LARP6 binding assay. **(F)** ITC curves of *COL1A1*-TIS binding to LARP6. **(G)** Dissociation constant (Kd) values of LARP6 binding to target RNAs; n.d, not detected.

We performed competitive binding experiments of labeled A1 with non-labeled LARP6 binding RNA targets (Figure **4B**). *COL1A1*-TIS showed a higher binding affinity to LARP6 (IC50, 2.71 nM) compared to A1 (IC50, 6.84 nM), suggesting that LARP6 has binding capacity that coordinates with mRNAs beyond 5’SL in *COL1A1* (Figure **4C-D**). LARP6 also showed binding with CDS of *COL5A1* and *LRP1*, indicating its interaction with mature mRNA structure (Figure **4C-D**). The RNA binding properties of LARP6 were also characterized in solution by ITC (Figure **4E**). Consistent with the TR-FRET results, ITC confirmed tight binding of LARP6 to *COL1A1-*TIS. The binding dissociation constant (Kd) for *COL1A1*-TIS was 0.4 μM, which is lower than the Kd value for *COL1A1*-5’SL (A1) (Figure **4F-G**). The titration of *LRP1*- and *COL5A1*-CDS showed direct binding of LARP6 to mature mRNA, with binding affinities in the low micromolar range (Figure **4F-G**, Figure **S6**). *CCNI* did not show binding in neither TR-FRET nor ITC, suggesting that interaction with LARP6 requires additional properties beyond *in-vitro* conditions.

### LARP6 enhances mRNA translation via direct interaction in their 5’UTR

We further evaluated the binding properties of LARP6 on mRNA. Interestingly, by plotting eCLIP fold-enrichment of immunoprecipitation (IP) over size-matched input (SMInput) of the binding sites according to the mature mRNA coordinates (i.e. 5’UTR, CDS and 3’UTR), a significant enrichment in binding is observed only within 5’UTRs, suggesting that LARP6 likely functions primarily as a translation regulator (Figures **5A-C** and Figures **S7A-C**).

**Figure 5.**
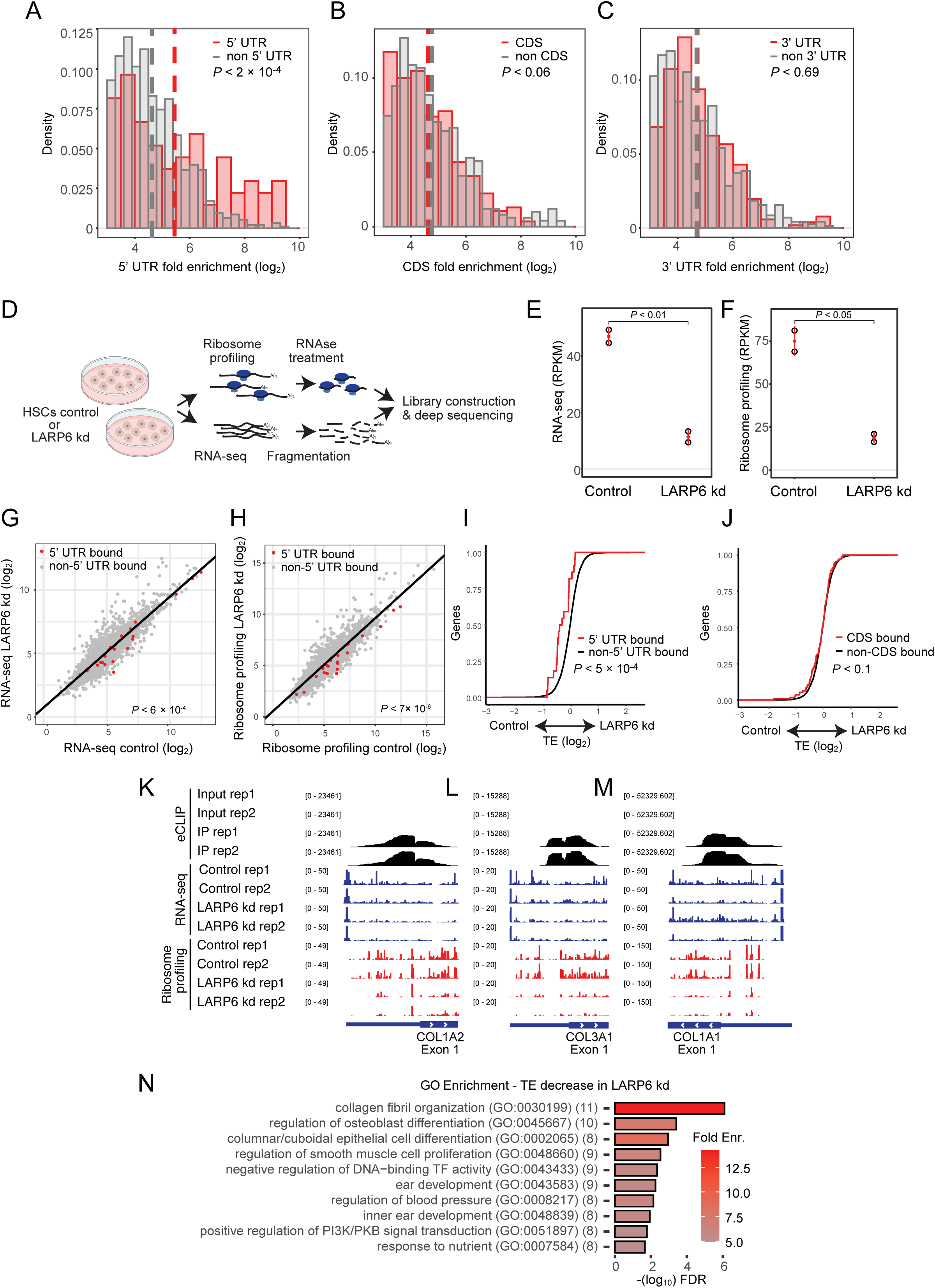
LARP6 directly regulates translation via binding to 5’UTRs. **(A-C)** Density analysis of fold enrichment (log2) from TGFβ1-stimulated HSC eCLIP analysis. **(A)** Non-5’UTR peaks and 5’UTR peaks are represented with grey and red bars, respectively, and the mean is indicated with a dashed line. The same analysis for CDS **(B)** and **(C)** 3’UTR targets. *P* value was calculated using student’s *t*-test. **(D)** Schematic illustration of ribosome profiling and RNA-seq in control and LARP6 knockdown HSCs. **(E-F)** LARP6 knockdown efficiency using LARP6-targeting dsiRNA in human HSCs (donor D22) indicated in **(E)** RNA-seq and **(F)** ribosome profiling data in RPKM. **(G-H)** Correlation of **(G)** RNA-seq and **(H)** ribosome profiling data in dsi- negative control (Control) and LARP6-targeting dsiRNA-transfected (LARP6 kd) HSCs. 5’UTR targets from eCLIP data are marked with red dots while non-bound targets are marked with grey dots. *P* value was calculated using student’s *t*-test. **(I-J)** Cumulative translation efficiency was calculated with the ratio of ribosome profiling to RNA-seq data in LARP6 knocked down cells to control. The **(I)** 5’UTR and **(J)** CDS targets from eCLIP data are marked with a red line, and the non-bound targets with a black line. *P* value was calculated using student’s *t*-test. **(K-M)** eCLIP, RNA-seq, and ribosome profiling read counts for **(K)** *COL1A2*, **(L)** *COL3A1*, and **(M)** *COL1A1* transcripts are represented in black, blue, and red, respectively. eCLIP data was generated in TGFβ1-treated HSCs. RNA-seq and ribosome profiling data were generated from control or LARP6 knocked down HSCs. **(N)** GO-enrichment analysis was calculated with negatively translationally regulated targets in LARP6 kd cells, using xtail analysis and FDR<0.1. Color represents fold enrichment, and the number of genes in each GO term is represented in parentheses.

To assess the role of LARP6 as a translation regulator, we conducted RNA-seq and ribosome profiling in LARP6-targeting dsiRNA and dsi-negatvie control-transfected human HSCs (Figures **5D-F**). We focused on genes that exhibit sufficient reads coverage (≥30) from both RNA-seq and ribosome profiling data, and significantly enriched in 5’UTR interaction from our CLIP data. We observed 22 targets from the (+)TGFβ1-stimulated HSCs eCLIP (Figures **5G-H**), and 14 targets from the (-)TGFβ1-stimulated HSCs (Figures **S7D-E**). We found that transcripts bound by LARP6 in their 5’UTR are more highly expressed and translated in control HSCs relative to LARP6 knockdown HSCs (Figures **5G-H** and **S7D-E**). Since mRNA translation reflects RNA abundance, we measured the translation efficiency (TE) in those conditions which measures the coverage of ribosomes over transcripts. We found a significant decrease in translation efficiency of 5’UTR LARP6 targets in LARP6 knockdown HSCs (Figure **5I** and Figure **S7F**), whereas the TE of CDS LARP6 targets show no change (Figure **5J** and Figure **S7G**), indicating that LARP6 binding to 5’UTRs directly enhance translation of mRNAs. As expected, *COL1A2*, *COL3A1* and *COL1A1* were found to be translationally regulated in ±TGFβ1 conditions, and demonstrate a significant reduction in translation efficiency with knockdown of *LARP6* (Figures **5K-M**). Finally, we examined enriched pathways that are regulated by LARP6 at the translational level. We calculated the significant changes in TE using xtail^26^ and found 171 genes that show a decrease in translation efficiency (TE) with LARP6 knockdown HSCs (Table **S2**, 163 genes show increase in TE. FDR≤ 0.1). The most significantly suppressed pathway in translation in *LARP6* knockdown HSCs was collagen fibril organization (Figure **5N**), similar to the enriched pathways from the eCLIP analysis (Figure **3C**). These data indicate the main role of LARP6 as a positive mRNA translation regulator of collagen-related pathways.

### LARP6 is necessary and sufficient for regulating translation

We aimed to assess the effect of LARP6 on reporter expression by cloning 5’UTRs of genes that are both bound by LARP6 in our eCLIP data and exhibit a significant decrease in TE with LARP6 depletion. This includes 5’UTRs of the collagen mRNAs *COL1A2*, *COL1A1*, *COL3A1* and *CCDC85B*, *SPTBP1* and *MAP4K4*. To evaluate if their respective 5’UTR sequences are sufficient to reflect LARP6 regulation, we cloned these upstream of firefly luciferase and expressed them in HSCs in control and in LARP6 knockdown conditions (Figure **6A**, and Figure **S8A-B**). Indeed, we observed a reduction in luciferase expression with LARP6 knockdown in the 5’UTR constructs (Figure **6B**). We sought to examine the effect of collagen mRNA TIS structural elements (i.e. collagen structural elements) with reduction of LARP6 in the cells. For that purpose, we cloned the structural elements upstream to firefly luciferase with and without canonical ATG, for maintaining either coding potential or structural potential (Figure **S8A-C**). The collagen structural elements show no effect with reduction of LARP6 levels, suggesting that these reporter elements are less sensitive to low levels of LARP6 in naive wild type HSCs (Figure **6C**). To examine if increase in LARP6 is sufficient to regulate the constructs expression, we exogenously expressed LARP6 in HeLa cells, which express low levels of endogenous LARP6 (Figure **6D** and Figure **S8D-E**). We observed a consistent increase in reporter expression, suggesting that LARP6 is sufficient to stimulate expression of 5’UTR containing targets but may ultimately depend on other unique regulatory features of the mRNAs or HSCs trans-regulators (Figure **6E**). Strikingly, collagen structural elements increase reporter expression with exogenously expressed LARP6, indicating high sensitivity of cis-regulatory elements for downstream coding sequences. Interestingly, preserving the canonical ATG in COL1A1 structural element was the only element that enhance the reporter expression beyond the structural element without the canonical ATG, suggesting an orchestrated regulation of LARP6 with the translation initiation and elongation machineries for COL1A1 gene expression (Figure **6F**). Altogether, our data shows that LARP6 is necessary and sufficient to regulate translation by interaction with 5’UTRs of targets beyond the known collagen mRNAs and mediates gene expression through structural elements of collagen mRNAs flanking their canonical translation initiation site.

**Figure 6.**
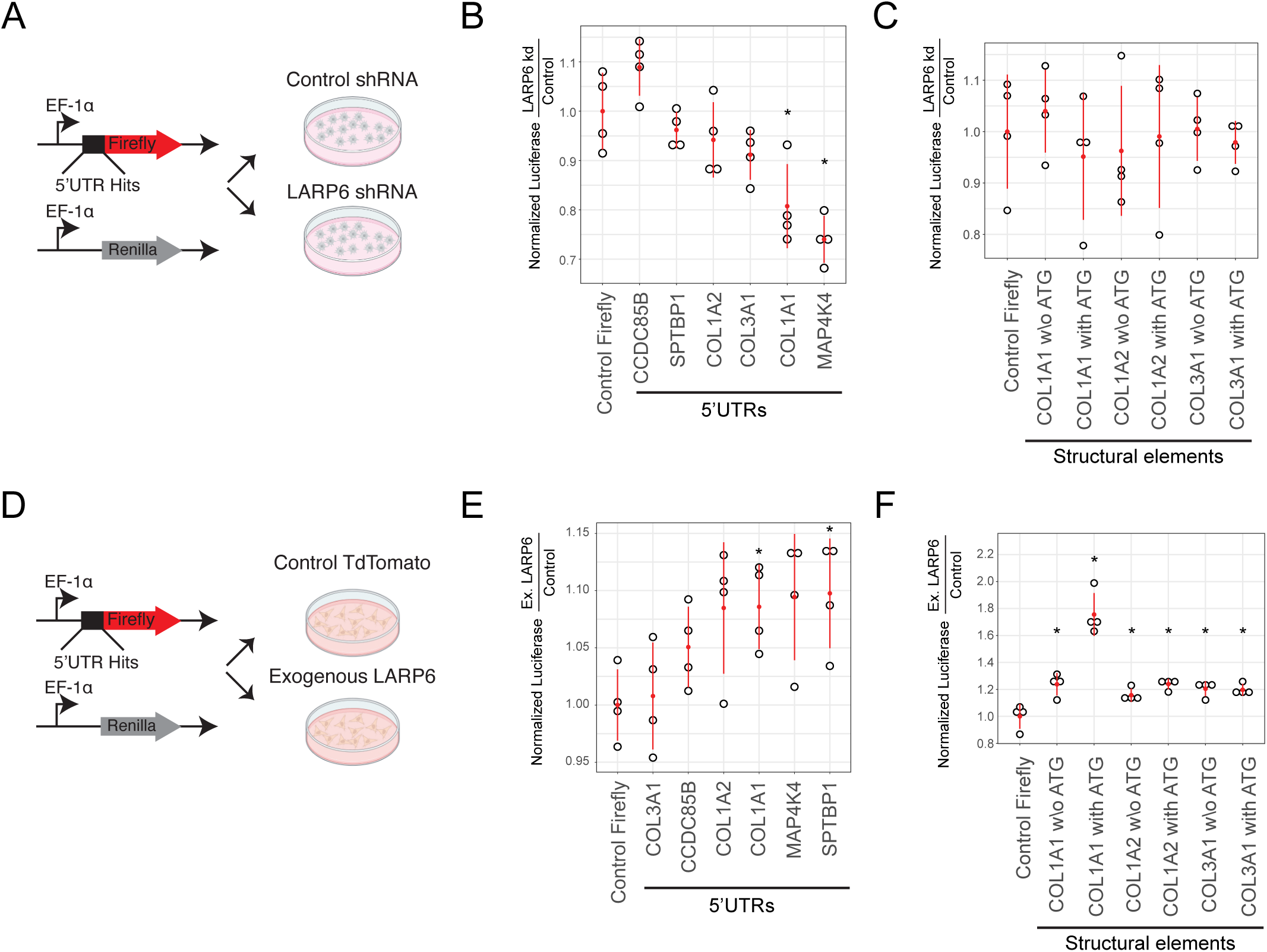
LARP6 regulates reporter expression with LARP6 5’UTR targets. **(A)** Schematic illustration of 5’UTRs cloning and transduction of control shRNA or LARP6 shRNA in human HSCs (donor D22). **(B-C)** Luciferase reporter fold change expression in LARP6 knocked down HSCs relative to control cells with reporter of **(B)** 5’UTR hits and **(C)** collagen structural elements with and without canonical ATG. Significant *P* < 0.05 is marked with an asterisk and calculated using student’s *t*-test. **(D)** Schematic illustration of 5’UTRs cloning and transduction in HeLa control or LARP6 exogenous expression in cells. **(E-F)** Luciferase reporter fold change expression in LARP6 exogenously expressed in HeLa cells relative to control cells with reporter of **(E)** 5’UTR hits and **(F)** collagen structural elements with and without canonical ATG. Significant *P* < 0.05 is marked with asterisk and calculated using student’s *t*-test.

### HSC-specific knockdown of LARP6 reduces liver fibrosis in human liver spheroids

Human liver spheroids serve as a useful tool to investigate the pathogenesis of metabolic liver diseases and are superior to 2D liver cell cultures or cultured liver slices^27^. To model the role of LARP6 in human liver fibrosis, we generated human liver spheroids that consists of human hepatocytes (donor D23), HSCs (donor D19), and other nonparenchymal cells (NPCs, donor D22) in the liver and can recapitulate liver fibrosis induced by metabolic stress (Figure **S2A**). Human liver spheroids were cultured in MASH- or MetALD-cocktail (MASH-cocktail supplemented with ethanol), mimicking metabolic injury complicated by chronic alcohol exposure (Figure **7A**)^28^. Both MASH- and MetALD-induced human liver spheroids exhibited significant lipid accumulation within the spheroids (Figure **7B**). In addition, exposure to ethanol accelerated expression of fibrogenic markers. Specifically, the expression of fibrogenic genes such as *COL1A1*, *COL1A2*, and *SERPINE1* increased in MetALD-induced spheroids compared to MASH-induced spheroids (Figure **7C**). Collagen Type I protein and CYP2E1 protein expression also exhibited a further increase in MetALD-induced spheroids (Figure **7D**).

**Figure 7.**
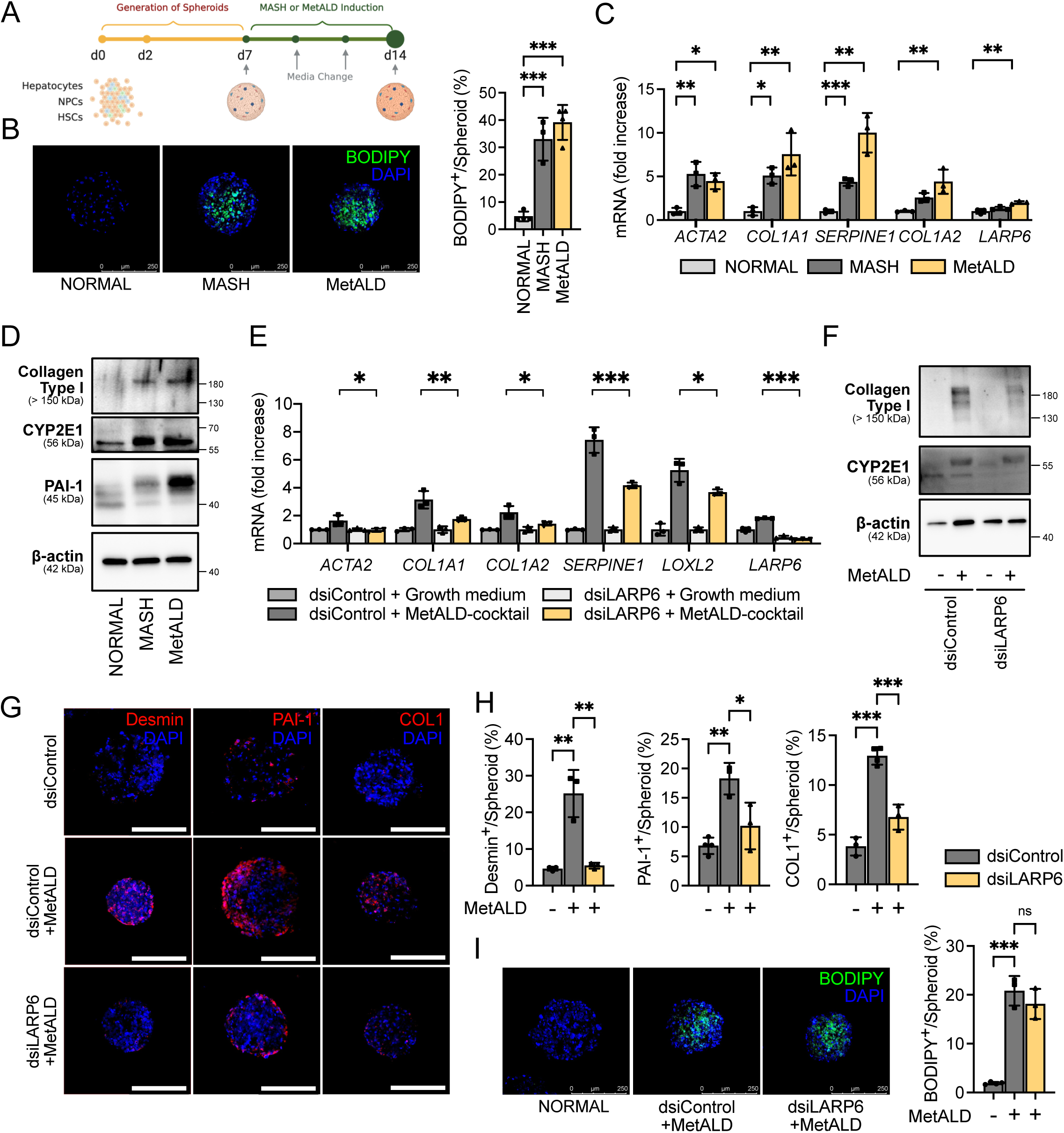
HSC-specific knockdown of LARP6 inhibits MetALD-induced fibrosis in human liver spheroids. **(A)** Schematic illustration of human liver spheroid generation. Human liver spheroids were generated using hepatocytes (donor D23), NPCs (donor D22), and HSCs (donor D19). **(B)** Immunofluorescent images of human liver spheroids stained with DAPI and BODIPY (scale bar = 250 µm). The BODIPY-positive area was normalized by spheroid area and calculated as percentage. **(C)** Expression of fibrogenic markers in MASH- or MetALD-induced human liver spheroids was evaluated using qRT-PCR analysis, and **(D)** Western blotting. **(E-I)** Human liver spheroids were generated using hepatocytes (donor D23), NPCs (donor D22), and LARP6 targeting dsiRNA-transfected HSCs (donor D19) and incubated under MetALD conditions. **(E)** Expression of fibrogenic markers in liver spheroids was assessed using qRT-PCR. **(F)** Western blotting was performed to measure Collagen Type I and CYP2E1 expression. **(G, H)** Human liver spheroids were stained for Desmin, PAI-1, Collagen Type I (COL), and DAPI (scale bar = 250 µm). Desmin, PAI-1, or COL1-positive area was normalized by spheroid area and calculated as a percentage. **(I)** Human liver spheroids generated with LARP6-targeting dsiRNA transfected HSCs were stained with DAPI and BODIPY (scale bar = 250 µm). The BODIPY-positive area was normalized by spheroid area and calculated as a percentage. Data are presented as mean ± SD; \**P* < 0.05, \*\**P* < 0.01, and \*\*\**P* < 0.001, one-way ANOVA followed by Tukey’s test.

Next, LARP6 protein was knocked down in human HSCs, which were then used to generate liver spheroids, followed by incubation with MASH or MetALD cocktails. HSC-specific knockdown of LARP6 significantly suppressed *ACTA2*, *COL1A1*, *COL1A2*, *SERPINE1*, and *LOXL2* mRNA expression in MetALD-induced spheroids (Figure **7E**). Knockdown of LARP6 led to a dramatic reduction in Collagen Type I expression in MetALD spheroids (Figure **7F-H**). Similarly, knockdown of LARP6 in HSCs reduced liver fibrosis induced by MASH cocktail (Figure **S9A-B**). These results support the idea that LARP6 regulates the translation of collagen-related mRNAs. HSC-specific knockdown of LARP6 did not affect lipid accumulation in the spheroids, suggesting that LARP6 plays a specific role in HSC activation rather than in hepatocytes (Figure **7I**).

## Discussion

Due to the rising prevalence of obesity and increased alcohol consumption, liver fibrosis associated with steatotic liver disease has increased substantially in the past decade^29^. Liver fibrosis occurs in response to chronic liver injury and is characterized by an excessive accumulation of extracellular matrix proteins^3^. Fibrillar collagens are the predominant ECM in human fibrotic liver tissues, and the development of liver fibrosis from F1 to F3 stages requires a progressive accumulation of Collagen Type I and III^2^.

LARP6, an RNA-binding protein with a specificity for fibrillar collagen mRNAs, is highly expressed in activated HSCs, thus presenting a promising therapeutic target for liver fibrosis. Previous studies demonstrated binding of LARP6 to a well-defined 5’SL structure in collagen mRNAs^25^. To examine LARP6 binding in HSCs, we applied eCLIP to capture RBP-RNA interactions transcriptome-wide^30^. Indeed, we found that LARP6 targets are highly represented by the collagen transcripts *COL1A1*, *COL1A2* and *COL3A1*, via direct interaction with a structural element that flanks their canonical TIS. These interactions were deployed through the bulge region of the mRNA stem. We also identify previously unknown LARP6 targets such as *CCNI* and *LRP1*. *CCNI*, which encodes the Cyclin I protein, is part of cell-cycle regulating proteins, which were found to be differentially expressed in HSCs during liver fibrosis^31^. Moreover, other Cyclin genes were found to promote HSCs proliferation during liver fibrosis^32^, suggesting that LARP6 may regulate HSCs proliferation during HSCs activation via interaction with *CCNI* 5’UTR. *LRP1,* which encodes the LDL Receptor Related Protein 1, has a controversial role in fibrosis. In rat kidney fibrosis model, *LRP1* activation promotes fibrosis development^33^. In activated HSCs, LRP1 signaling is regulated via interactions with connective tissue growth factors (CTGFs)^34^. However, LRP1 global knockout mice had higher levels of activated HSCs and αSMA signal, as well as collagen expression, which implies anti-fibrogenic activity^35^. In our data, we show a significant reduction in mRNA expression and translation of LRP1 mRNA in LARP6 depleted HSCs, suggesting that LARP6 enhances expression of *LRP1*, presumably to promote HSCs activation and fibrosis.

LARP6 binding sites were found in mature mRNAs, including 5’UTRs, CDSs and 3’UTRs, implying an extensive regulation by LARP6 as an RBP. We found that LARP6 binds to *LOXL2*^36^, *PLOD3*^37^, *COL5A1*, *COL5A2*^38,39^ and *COL18A1*^40^, all part of collagen fibril organization and extracellular matrix pathways and associated with fibrosis progression. Interestingly, some of these interactions reside on exons separated by intronic regions, suggesting that structural regulatory elements are formed only on mature mRNA for regulating mRNA fate in the cytoplasm (Figure **4**). In this work, we extend the LARP6 binding repertoire from the previous three collagen mRNAs to 397 genes and identify potential targets whose expression may be stimulated in fibrosis due to increased expression of LARP6.

LARP6 binding is highly enriched in the 5’UTR suggesting that LARP6 controls mRNA translation. We further explored the role of LARP6 as an mRNA translation regulator by employing ribosome profiling^41^ in human HSCs. By combining our eCLIP measurements for 5’UTR binding with ribosome profiling we found a total of 22 genes that are significantly translationally regulated in LARP6 knockdown HSCs, including collagen genes, demonstrating a direct regulation of translation by LARP6 binding. The structural elements of collagen genes overlap the TIS, suggesting that LARP6 binds to these elements and promotes translation, likely via recruitment of RNA helicase to enhance translation. Such a helicase, RNA helicase A, was shown to be expressed mainly in activated HSCs, and was tethered to the 5’SL structure of collagen genes through recruitment by the C-terminal domain of LARP6^42^. mRNA translation assays revealed a new set of 334 targets that exhibit translational dependency on LARP6 expression. With the genes that exhibit a decrease in TE with LARP6 depletion (171 genes, 0.1>FDR) we captured 168 novel genes beyond *COL1A1*, *COL1A2* and *COL3A1*. These genes include *SLC16A1* (MCT1), which haplosufficiency in mice increased resistance to hepatic steatosis development^43,44^*, AOX1,* which was found to alleviate liver fibrosis with decrease in expression^45^. Our results demonstrate for the first time the complete set of targets of LARP6 on cellular mRNA expression and translation in HSCs, revealing that while collagen genes are the most enriched targets for translation, LARP6 binds an extensive list of transcripts providing novel targets that are regulated at the post-transcriptional level.

Knockdown of LARP6 in human HSCs decreased not only collagen expression, but also other fibrogenic markers such as αSMA and PAI-1. It is possible that decreased collagen expression regulates TGFβ1-induced HSC activation. HSCs are known to express two types of collagen receptors: integrins and discoidin domain-containing receptors (DDRs)^46^. Integrin-dependent interaction with the ECM promotes a profibrogenic phenotype of activated HSCs^47^. Endogenous collagen synthesis induces DDR2 expression and phosphorylates DDR2, thereby promoting cell proliferation and differentiation^48,49^. DDR2 was indeed identified as one of the collagen fibril organization genes depleted in our ribosome profiling of LARP6 knockdown human HSCs. This result suggests the possibility that reduction in collagen synthesis caused by LARP6 depletion regulates the activation of human HSCs via downregulation of DDR2.

To date, there are no therapies targeting liver fibrosis. Advances in drug development for liver fibrosis have been limited by *ex vivo* models that fail to fully recapitulate the complex clinical, histological, and molecular features of metabolic liver diseases^27^. Here, we demonstrate the physiological role of LARP6 on HSC activation using human HSCs isolated from MASH or MetALD livers. Moreover, we examined the effect of HSC-specific knockdown of LARP6 on liver fibrosis induced by metabolic stress in human liver spheroids. Unlike most human liver organoid systems composed primarily of hepatocyte-like cells, which may not fully address the multicellular aspects of metabolic liver diseases^50^, our human liver spheroid system mimics liver fibrosis induced by metabolic injuries using all cell types present in the liver. LARP6 selectively targets collagen subtypes involved in liver fibrosis and is a promising therapeutic approach for human liver fibrosis from various etiologies. The significance of LARP6 has led to initial attempts to develop an inhibitor^5^, and further investigation is necessary to develop a lead compound that can efficiently inhibit LARP6 binding to collagen mRNAs.

In conclusion, we demonstrate the role and mechanism of LARP6 in stimulating human liver fibrosis. Depletion of LARP6 in human HSCs decreases fibrogenic gene expression and translation, which directly reduces collagen biosynthesis via binding to collagen mRNAs, and via extensive interactions with cellular mRNA. HSC-specific knockdown of LARP6 inhibited the development and progression of liver fibrosis induced by metabolic stress in human liver spheroids. Based on these results, targeting LARP6 in human HSC may become a novel strategy to treat metabolic dysfunction-associated liver fibrosis.

## Acknowledgements

This research was supported by the Martha Proctor Mack Foundation, by the National Institutes of Health HG009889, HG004659, HG011864 and CA273432 to GWY, R01DK111866, R56DK088837, DK099205, AA028550, DK101737, AA011999, DK120515, AA029019, DK091183, P42ES010337, R44DK115242 (DAB and TK), R01CA285997 (DAB), by Sanford Stem Cell Innovation Center (GWY) and Sanford Stem Cell Fitness and Space Medicine Center at Sanford Stem Cell Institute (UCSD) (TK). The shared resources at SBP were supported by the National Cancer Institute Grant P30 CA030199. Work at the Center for Epigenomics was supported in part by the UC San Diego School of Medicine. This work was partially supported by the National Institutes of Health, Grant 2UL1TR001442 of CTSA.

We thank Lifesharing OPO and UCSD microscopy core (NINDS P30NS047101) for their support. We thank Karin Diggle for her excellent technical support. HYK was supported by a basic science research program through the National Research Foundation of Korea funded by the Ministry of Education RS-2023-00245179. OM was supported by the Gruss-Lipper postdoctoral fellowship. The authors used BioRender.com to create the illustrations.

## Author contributions

HYK, OM, conceptualization, investigation, visualization, writing-original draft, writing-review & editing; WL, methodology, investigation; SBR, formal analysis; CH, BAY, SMB, JD, JJ, KH, HJ, C-TM, AB, investigation; ES, MJ, BS, conceptualization; TK, funding acquisition, supervision, writing-review & editing; GWY, conceptualization, funding acquisition, supervision, writing-original draft, writing-review & editing; DAB, conceptualization, funding acquisition, supervision, writing-original draft, writing-review & editing.

## STAR Methods

**KEY RESOURCES TABLE**

| REAGENT or RESOURCE | SOURCE | IDENTIFIER |
| --- | --- | --- |
| **Antibodies** |  |  |
| Rabbit anti- αSMA antibody | Abcam | ab5694 |
| Rabbit anti-Collagen type I antibody | Rockland | 600-401-103S |
| COL1A1 (E8F4L) XP<sup>®</sup> Rabbit monoclonal antibody | Cell Signaling | 72026s |
| Mouse monoclonal anti-β-actin antibody | Sigma | A5441 |
| Rabbit anti-PAI-1 antibody | Proteintech | 13801-1-AP |
| Mouse anti-LARP6 polyclonal antibody | Novus Biologicals | H00055323-B01P |
| Rabbit anti-LARP6 polyclonal antibody | Sigma | HPA049029 |
| Rabbit anti-Vimentin polyclonal antibody | Proteintech | 10366-1-AP |
| Mouse anti-HNF4α monoclonal antibody (K9218) | Invitrogen | MA1-199 |
| Mouse anti-Desmin monoclonal antibody | Proteintech | 60226-1-Ig |
| Mouse anti-GAPDH monoclonal antibody, clone 6C5 | Sigma-Aldrich | MAB374 |
| Rabbit anti-CYP2E1 polyclonal antibody | Abcam | ab1252 |
| CD68 (D4B9C) XP<sup>®</sup> Rabbit monoclonal Ab | Cell Signaling | 76437s |
| Goat anti-Rabbit IgG, HRP | Invitrogen | 31460 |
| Goat anti-Mouse IgG HRP | Invitrogen | 31430 |
| Goat anti-Mouse IgG (H+L) Cross-Adsorbed Secondary Antibody, Alexa Fluor<sup>™</sup> 488 | Invitrogen | A-11001 |
| Donkey anti-Rabbit IgG (H+L) Highly Cross-Adsorbed Secondary Antibody, Alexa Fluor 594 | Invitrogen | A-21207 |
| **Biological samples** |  |  |
| Human livers | Lifesharing OPO | D1-D23<br>See Figure **S1A** and Figure **S2A** |
| **Chemicals, peptides, and recombinant proteins** |  |  |
| Nycodenz | Axell | AN1002424 |
| AOF BP Protease | VitaCyte | 003-1000 |
| Collagenase MA | VitaCyte | 001-2030 |
| Dulbecco's-modified Eagle's medium (DMEM) | Gibco | 11965-092 |
| Percoll | Cytiva | 17089101 |
| Fetal Bovine Serum (FBS) | Gemini Bio | 100-106 |
| Penicillin-Streptomycin (PS) | Gibco | 10378-016 |
| Dexamethasone | Invivogen | tlrl-dex |
| Insulin-transferrin-selenium | Gibco | 41400-045 |
| HEPES | VWR | 45000-690 |
| 10× DPBS | VWR | 45000-426 |
| Antibiotic-antimycotic | Gibco | 15240062 |
| Trypsin-EDTA (0.25%), phenol red | Gibco | 25200056 |
| 1× PBS, without calcium and magnesium | Corning | 21-040-CV |
| Insulin-Transferrin-Selenium (ITS -G) (100×) | Thermo | 41400045 |
| Palmitate | Sigma | P0500 |
| Oleate | Sigma | O1257 |
| Fructose | VWR | 97061-236 |
| Lipopolysaccharides (LPS) | Invivogen | tlrl-3pelps |
| Ethanol | Koptec | V1016 |
| Bovine serum albumin solution | Sigma | A1595 |
| D-(+)-Glucose solution | Sigma | G8644 |
| Recombinant Human TGF-β1 | Proteintech | HZ-1131 |
| RIPA lysis buffer | Teknova | R3792 |
| Protease and Phosphatase Inhibitor Cocktail | Thermo | 78443 |
| 4% paraformaldehyde | Thermo | 28906 |
| Hematoxylin Counterstain | VectorLabs | H-3401-500 |
| Lipofectamine™ RNAiMAX Transfection Reagent | Invitrogen | 13778075 |
| Triton™X-100 | Sigma | T9284 |
| Horse Serum | Gibco | 26050-088 |
| Antifade Mounting Medium with DAPI | VECTASHIELD | H-2000 |
| Tissue-Tek® O.C.T. Compound | Sakura | 4583 |
| BSA Fraction V | Sigma | 03116956001 |
| Sucrose | Sigma | S0389 |
| BODIPY™ 493/503 | Thermo | D3922 |
| Fast SYBR™ Green Master Mix | Life Technologies | 4385617 |
| Super Signal™ West Pico PLUS Chemiluminescent Substrate | Thermo | 34580 |
| RNase I | Invitrogen | AM2295 |
| anti-rabbit IgG Dynabeads | Thermo | 11204D |
| FastAP Thermosensitive Alkaline Phosphatase | Thermo | EF0651 |
| T4 Polynucleotide Kinase | NEB | M0201S |
| T4 RNA ligase | NEB | M0437M |
| Proteinase K | NEB | P8107S |
| SUPERase·In™ RNase Inhibitor | Thermo | AM2694 |
| TRIzol | Invitrogen | 15596-018 |
| DNase I | NEB | M0303S |
| SuperScript™ III First-Strand Synthesis SuperMix | Invitrogen | 18080400 |
| Critical commercial assays |  |  |
| RNeasy® Plus Micro Kit (50) | Qiagen | 74034 |
| DAB Substrate Kit, Peroxidase (HRP) | Vector Laboratories | SK-4100 |
| High-Capacity cDNA Reverse Transcription Kit | Life Technologies | 4368813 |
| Pierce™ BCA Protein Assay Kit | Thermo | 23225 |
| Dual-Luciferase® Reporter Assay System | Promega | E1910 |
| Dynabeads™ mRNA DIRECT™ Purification Kit | Invitrogen | 61012 |
| Deposited data |  |  |
| GSE244832<sup>15</sup> | https://www.ncbi.nlm.nih.gov/geo/query/acc.cgi?acc=GSE244832 | GEO: GSE244832 |
| TF-target gene database<sup>21</sup> | Plaisier et al.<sup>21</sup> | http://tfbsdb.systemsbiology.net |
| GSE279065 | https://www.ncbi.nlm.nih.gov/geo/query/acc.cgi?acc=GSE279065 | GEO: GSE279065 |
| Experimental models: Cell lines |  |  |
| HeLa cells | ATCC | CCL-2 |
| Oligonucleotides |  |  |
| Primers for qPCR | IDT | Table **S3** |
| RNA oligos for FRET and ITC analysis | IDT | Table **S3** |
| Reporter element | IDT | Table **S3** |
| non-target shRNA SHC002 | Sigma | MISSION shRNA library |
| shRNA for knocking down LARP6 | Sigma | Table **S3**; MISSION shRNA library, TRCN0000145388 |
| Predesigned DsiRNA | IDT | hs.Ri.LARP6.13.1 |
| Predesigned DsiRNA | IDT | hs.Ri.LARP6.13.2 |
| Predesigned DsiRNA | IDT | hs.Ri.LARP6.13.3 |
| Negative Control DsiRNA | IDT | 51-01-14-04 |
| Software and algorithms |  |  |
| GraphPad Prism (v10.0) | GraphPad Prism | GraphPad Prism |
| Leica Application Suite X (LAS X) | Leica Application Suite X (LAS X) | Leica Application Suite X (LAS X) |
| QuPath (v0.5.0-rc1) | QuPath | QuPath |
| ImageJ (v2.3.0) | ImageJ | ImageJ |
| Suerat (v4.0) | Satija lab | https://satijalab.org/suerat/ |
| NanoAnalyze software | TA Instrument | TA Instrument |
| Bowtie v1.1.2 | Langmead et al.<sup>52</sup> | https://genomebiology.biomedcentral.com/articles/10.1186/gb-2009-10-3-r25 |
| Xtail | Xiao et al.<sup>26</sup> | https://www.nature.com/articles/ncomms11194 |
| eCLIP analysis pipeline CLIPper | Blue et al.<sup>53</sup> | https://doi.org/10.5281/zenodo.5076591 |
| IDR (2.0.2) | Li et al.<sup>54</sup> | https://projecteuclid.org/journals/annals-of-applied-statistics/volume-5/issue-3/Measuring-reproducibility-of-high-throughput-experiments/10.1214/11-AOAS466.full |
| LocARNA | Lorenz et al.<sup>55</sup><br>Will et al.<sup>56</sup> | http://www.bioinf.uni-freiburg.de/Software/LocARNA/ |
| RNAfold 2.3.1 | Lorenz et al.<sup>55</sup> | http://rna.tbi.univie.ac.at/ |
| STAR (2.7.6a) | Dobin et al.<sup>57</sup> | https://pubmed.ncbi.nlm.nih.gov/23104886/ |
| Other |  |  |
| 96-well clear round bottom ultra-low attachment surface spheroid microplate | Corning | 4520 |
| Immobilon®-P transfer membrane | Millipore | IPVH0010 |
| 1536-well assay plate | Corning | 3725 |

## RESOURCE AVAILABILITY

### Lead contact

Further information and requests for resources and reagents should be directed to and will be fulfilled by the lead contact, David A. Brenner.

### Materials availability

- This study did not generate new unique reagents.

### Data and code availability

- Data accession number reported in this paper is GEO: GSE279065. Note to reviewers: the data will be made publicly available upon publication of this manuscript. You can access the data using the access token cjkjyoqubpizdoh

at https://www.ncbi.nlm.nih.gov/geo/query/acc.cgi?&acc=GSE279065.

- Any additional data reported in this paper are available from the lead contact upon request.

## EXPERIMENTAL MODEL AND STUDY PARTICIPANT DETAILS

### Human livers

Deidentified donor livers (IRB 171883XX), certified as “no human subjects” (according to the Code of Federal Regulations Title 45, part 46 and UCSD Standard Operating Policies and Procedures) by HRPP Director and IRB Chair, were obtained through Lifesharing organ procurement organization. Lifesharing provided the informed consent, laboratory tests (ALT, AST, liver biopsy, serology, and others), as well as patient’s history (alcohol consumption, cause of death, age, BMI, and gender). Livers were graded by a pathologist using a double-blinded method and identified as NORMAL, MASL, MASH, and MetALD (see Figure **S1A**, and **S2A**).

### Primary human HSC isolation and culture

Human HSCs were isolated from human liver tissues using collagenase/protease perfusion and gradient centrifugation with 8.2% Nycodenz^58^. Isolated human HSCs were amplified for at least 3 weeks prior to cryopreservation. Human HSCs in passage 2 or 3 were used for further experiments. Human HSCs were cultured in DMEM supplemented with 10% FBS and 1% antimycotic-antibiotic, at 37°C in 5% CO2.

## METHOD DETAILS

### Generation of human liver spheroids

Human liver spheroids were generated according to protocol^28^. Briefly, cryopreserved human hepatocytes were thawed and dead cells were removed by gradient-centrifugation (15 ml of 90% Percoll and 35 ml DMEM). Hepatocytes (3×10^5^), NPCs (1.5×10^5^), and HSCs (0.8×10^5^) were cultured in a 96-well clear round bottom ultra-low attachment surface spheroid microplate in a growth factor-enriched medium, which is DMEM supplemented with 10% FBS, 1% PS, 0.1 μM dexamethasone, and 1% insulin-transferrin-selenium. Spheroid microplates were shaken (450 rpm, 25 min), and the growth medium was exchanged after 2 days. Spheroids were generated after 7 days and induced with metabolic liver injury by incubation with MASH- or MetALD-cocktail for an additional 7 days. The MASH-cocktail contains 160 μM palmitate, 160 μM oleate, 10 mM fructose, 5.5 mM glucose, 2 μg/ml LPS, and 1 ng/ml human TGFβ1. The MetALD-cocktail contains 100 mM ethanol in addition to the MASH-cocktail.

### HSC-specific gene knockdown

Human HSCs (3.0 × 10^5^ per 10 cm dish) were transfected with dsiControl (dsi-negative control) or LARP6 targeting dsiRNA by incubation with RNAiMAX for 48 h according to the manufacturer’s protocol. Three different dsiRNA duplexes were tested, and the one with the highest knockdown efficiency (hs.Ri.LARP6.13.3), was used for further experiments. A dsiRNA negative control was transfected into human HSCs as a control.

### Histological analysis

Paraffin embedded liver sections were stained for H&E, Masson’s trichrome, Sirius Red, Desmin, αSMA, CD68, LARP6, and CYP2E1, followed by DAB staining. Brightfield images were captured using Olympus Microscope (IX71), and stained areas were measured using QuPath software.

### Immunofluorescent staining of human HSCs

Human HSCs were fixed with 4% paraformaldehyde for 30 min, permeabilized with 0.2% Triton™X-100, followed by incubation with 10% horse serum for 1 h and a rabbit anti-Collagen type I antibody for 16 h. HSCs were stained with Alexa Fluor 594-conjugated secondary antibody and mounted with an antifade mounting medium with DAPI. Immunofluorescent images were captured using Leica SP8 confocal microscopy and fluorescent-positive area were measured using QuPath software.

### Immunohistochemistry of human liver spheroids

Human liver spheroids were harvested and fixed with 4% paraformaldehyde for 30 min, followed by incubation in 30% sucrose for 2 days. Spheroids were embedded in OCT and cryosectioned to 10 μm. Cryosections were permeabilized with 0.2% Triton™X-100, blocked with 1% BSA, and stained for Collagen Type I, PAI-1, Desmin, Vimentin, and HNF4α. Neutral lipids were stained with 10 μg/ml BODIPY^TM^ 493/503 dye. Immunofluorescent images were captured using Leica SP8 confocal microscopy. Spheroid area and fluorescent-positive area were measured using QuPath software.

### SnRNA-seq quality control and integration

SnRNA-seq from 18 human liver samples were combined using Seurat’s data integration protocol^59^. Before filtering there were 150,656 nuclei. Cells were removed from further analysis if they had fewer than 200 features, greater than 7500 features, or greater than 5 percent mitochondrial reads. Following this filtering step, we retained 142,469 nuclei. Potential doublets were identified and removed using DoubletFinder. Ambient RNAs were removed by running CellBender, and additional genes from background contamination were filtered^60^. Following doublet filtering, we retained 78,184 nuclei to carry forward in the analysis. Nebulosa was used to display smoothed expression in the UMAP figures^51^.

### SnATAC-seq quality control and integration

SnATAC-seq from 18 human liver samples were combined using Signac’s data integration protocol^59^. Before filtering there were 101,335 nuclei. Doublets were removed using Scrublet prior to integration^61^. Further quality control filters were applied. Specifically, nuclei were retained in the analysis if they had greater than 500 fragments, less than 20,000 fragments, FRiP greater than 0.2, blacklist fraction less than 0.05, nucleosome signal less than 4, and TSS per cell greater than 7. Following doublet filtering, we retained 125,241 nuclei were retained to carry forward in the analysis.

### Quantitative Real-time polymerase chain reaction (qRT-PCR)

RNA was extracted from human HSCs or spheroids (16 spheroids per sample) with RNeasy^®^ Plus Micro kit, and reverse transcribed using High-Capacity cDNA reverse transcription kit. mRNA expression was assessed using Fast SYBR Master Mix and primers (Table **S3**).

### Western blotting

Human HSCs or spheroids (48 spheroids per sample) were lysed using RIPA lysis buffer supplemented with Halt™ Protease and Phosphatase Inhibitor Cocktail. Cell lysates were centrifuged at 16,000 g for 15 min, and protein content was measured with Pierce™ BCA protein assay kit. Protein samples were separated with sulfate-polyacrylamide gel electrophoresis and transferred Immobilon^®^-P transfer membrane. Immunoblots were incubated with primary antibodies followed by HRP-conjugated secondary antibody and protein expression was detected using chemiluminescent HRP substrate. Densitometric analysis was performed using ImageJ software.

### eCLIP

eCLIP on human HSCs was performed in two replicates as previously described^53^. Briefly, 2.0 × 10^7^ HSCs were treated with human TGFβ1 or remained untreated, then crosslinked, lysed (50 mM Tris-HCl pH 7.4, 100 mM NaCl, 1% (v/v) Igepal CA-630, 0.1% (v/v) SDS and 0.5% (wt/v) sodium deoxycholate) and sonicated. Lysate was treated with RNase I for RNA fragmentation. Anti-LARP6 antibody (Sigma) was pre-incubated with anti-rabbit IgG Dynabeads for 1 h at room temperature, added to lysate and incubated overnight at 4°C. For the input sample, 2% of each sample was kept separately. The immunoprecipitation (IP) sample was washed, and the RNA was dephosphorylated with FastAP, and T4 polynucleotide Kinase, followed by 3′ RNA adaptor ligation with T4 RNA ligase. 10% of IP and input samples were used for protein gel visualization for size indication and successful IP. IP and input samples were run on a protein gel and transferred to nitrocellulose membranes. Protein bands above the LARP6 protein size were excised from the membrane, followed by treatment with proteinase K for RNA release. Input samples followed the same treatment. The extracted RNA sample was reverse transcribed and performed PCR for library construction.

### eCLIP Data Analysis

eCLIP reads were processed as previously described^53^. Briefly, unique molecular identifiers (UMIs) were extracted with UMI tools (1.0.0), followed by trimming reads with cutadapt (2.5). Processed reads were mapped to repeat elements (RepBase) database with STAR (2.7.6a) for rRNA and repeat elements filtering, and the remaining reads to GRCh38. Uniquely mapped reads were sorted, deduplicated, and used with CLIPper^53^ (available at https://github.com/YeoLab/clipper) to call un-normalized peak clusters. Aligned reads of IP samples were compared to their size-matched input to produce normalized enriched peaks using scripts (overlap_peakfi_with_bam.pl and compress_l2foldenrpeakfi_for_replicate_overlapping_bedformat.pl) that are available at https://github.com/yeolab/eclip. Reproducibility of eCLIP peaks were calculated by ranking the normalized peaks in each replicate according to entropy values and irreproducibility discovery rate (IDR (2.0.2)) was used to generate a final list of reproducible eCLIP peaks. All pipeline definition files and scripts used to merge replicates are available at https://github.com/yeolab/merge_peaks.

### Multiple sequencing alignment for collagen mRNA

CLUSTAL W (1.81) multiple sequence alignment of 43 eutherian mammals EPO from ensemble was downloaded for hg38 genome, and structural region was selected based on structural region in figure 3E (without --Ancestral_sequences–). Then, structural minimum free energy (MFE) and structural conservation index (SCI) was calculated using the ViennaRNA package using RNAalifold by the following command: RNAalifold -p --MEA --sci --aln --color

### Isothermal titration calorimetry (ITC)

ITC was performed in Sanford Burnham Prebys Protein Production and Analysis Facility, using Low Volume Affinity ITC calorimeter (TA Instruments). 6 ml aliquots of solution containing between 30 and 50 mM ligand were injected into the cell containing 30 to 63 mM protein. 20 injections were made. The experiments were performed at 25°C in buffer containing 20 mM Hepes pH 7.5, 200 mM NaCl, 5 mM MgCl2, 10 mM β-mercaptoethanol. Baseline control data were collected injecting ligand into the cell containing the buffer only. ITC data were analyzed using NanoAnalyze software provided by TA Instruments. RNA oligo sequences used in this analysis are listed in Table **S3**.

### TR-FRET analysis

Assay buffer containing 25 mM HEPES pH 7.5, 100 mM NaCl, 5 mM MgCl2, and 0.005% Tween 20 was prepared fresh from concentrated stock the day of the experiment. 10 nM DIG-labeled A1 RNA was pre-mixed with various amount of competing unlabeled RNA (0-500 nM) and then 2ul was added to the 1536-well assay plate. 2 μl of 4.5 nM LARP6 (70-295aa) was added, and the mixture was incubated for one hour at room temperature. 0.5 nM Eu-anti-FLAG W1024 and 5 nM AF647-anti-DIG antibody was pre-mixed and 2 μl was added for additional one-hour incubation at room temperature. The plate was read on PHERAStar plate reader with HTRF^®^ 337/620/665 TR-FRET module. All concentrations were final assay concentrations after addition of all reagents to the assay plate. RNA oligo sequences used in this analysis are listed in Table **S3**.

### Ribosome profiling and RNA-seq libraries

dsiControl- or dsiLARP6-transfected human HSCs were harvested from a replicate of 10cm plates for each condition for ribosome profiling libraries as previously described^62^. Briefly, cells were treated with 100 μg/mL cycloheximide for 1 min at 37°C followed by washing with ice-cold PBS supplemented with 100 μg/mL cycloheximide. Cells were lysed with lysis buffer (12.5 mM Tris, pH 7.0; 7.5 mM Tris, pH 8.0; 150 mM NaCl; 5 mM MgCl2; 100 μg/mL cycloheximide; 1 mM DTT; 1% (v/v) Triton X-100; 20 U/mL DNase), collected and centrifuged at 16,000 g for 10 min at 4 °C. Cell lysate was then treated with 250 U RNase I at 25 °C for 45 min, followed by 200U SUPERase·In™ RNase Inhibitor for quenching. For pellet ribosome footprints, samples were loaded on sucrose cushion (34% sucrose, 20 mM Tris pH 7.5, 150 mM NaCl, 5 mM MgCl2, 1 mM dithiothreitol and 100 μg/ml CHX) and spin for 1 h at 100,000 rpm in a TLA-110 rotor (Beckman) at 4 °C. Then, the polysome pellet was resuspended in 1 mL TRIzol (15596-018, Invitrogen), and RNA was extracted by chloroform-based separation according to manufacturer instructions. 10μg of total RNA was used to run on a 15% TBE-UREA gel, and 28-34 mRNA fragments were extracted followed by ribosome profiling library construction as described^62^. Corresponding samples were grown in parallel for RNA-seq in 6 wells plate and treated as described^63^. Briefly, cells were collected with TRIzol, and RNA was extracted using manufacturer protocol. Then, RNA was used for poly A selection with Dynabeads mRNA DIRECT Purification Kit. mRNA was treated with DNA degradation and 3’ mRNA was resolved and phosphorylated with DNase I (M0303S, NEB) and 3′ dephosphorylation using FastAP Thermosensitive Alkaline Phosphatase and T4 polynucleotide kinase. The RNA was ligated with 3′ adaptor ligated using T4 ligase, and reverse transcribed with SuperScript™ III First-Strand Synthesis SuperMix for first-strand cDNA synthesis. cDNA products were ligated for second adaptor using T4 ligase and amplified by PCR for final library products^63^.

### Computational analysis for Ribosome profiling and RNA-seq

Sequencing reads were aligned as previously described^64^. In brief, linker 1 IDT (CTGTAGGCACCATCAAT) and poly(A) sequences were removed, and the remaining reads were aligned to rRNA. The unaligned rRNA reads were used for further alignment to GRCh38 human sequence. Alignment was performed using Bowtie v.1.1.229 with a maximum of two mismatches per read. Then, unaligned reads to the genome were used for alignment to sequences that spanned splice junctions. In ribosome profiling library analysis, the ribosome P site was calculated using the 5′ end of the reads of the ORFs with +12 for reads that were 28–29 bp and +13 for reads that were 30–33 bp.

### LocARNA structure and sequence alignment

Pairwise and multiple sequence alignment was performed using the online LocARNA software: http://www.bioinf.uni-freiburg.de/Software/LocARNA/. RNA structure prediction was performed using the Vienna RNA secondary structure server^65^.

### Reporter assay constructs

shRNA knockdown was performed using shRNA plasmid with the oligo in Table **S3**. Control plasmid was the non-target shRNA. For over-expression of LARP6, transcript variant 1 was expressed relative to tdTomato reporter. Sequences used for reporter elements are listed in Table **S3**.

### Transduction of shRNA into HSCs and HeLa cells

Lentivirus was generated for LARP6 short hairpin RNA (shRNA), control shRNA and the other vectors using transduction in HEK293T cells for 48 h. After transduction, LARP6 shRNA and control shRNA were transduced into HSCs with 10 μg/mL polybrene for 72 h and selected with puromycin. Then, cells were harvested and seeded in a 96 well-plate for dual luciferase reporter assay system. Different 5’UTR constructs were cloned upstream to firefly luciferase, while Renilla luciferase was used for normalization and transduced with 10 μg/mL polybrene for 24 h, followed by luminescence assay for luciferase signal. Parallel plates were used for measurement of LARP6 expression in the cells.

## QUANTIFICATION AND STATISTICAL ANALYSIS

Data are presented as mean ± S.D. Statistically significant differences were assessed using the unpaired Student’s *t*-test or one-way ANOVA followed by Tukey’s multiple comparison test. GraphPad PRISM software was used for the statistical analysis. *P* value < 0.05 was considered statistically significant.

### Supplemental information

Document S1. Figures S1-S9 and Table S3

**Table S1.** Excel file containing eCLIP significant windows in TGFβ1 treated HSCs and untreated HSCs, related to Figure 3

**Table S2.** Excel file containing translation efficiency measurements ribosome profiling data, related to Figure 5

