## Supporting Documents for "LARP6 regulates the mRNA translation of fibrogenic genes in liver fibrosis"

### Supporting Document for
### LARP6 regulates the mRNA translation of fibrogenic genes in liver fibrosis

Hyun Young Kim,[1,2,11] Orel Mizrahi,[3,4,5,11] Wonseok Lee,[1,2] Sara B. Rosenthal,[6] Cuijuan Han,[1] Brian A. Yee,[3,4,5] Steven M. Blue,[3,4,5] Jesiel Diaz,[3,4,5] Jyotiprakash Jonnalagadda,[3,4,5] Kanani Hokutan,[1,2] Haeum Jang,[1,2] Chen-Ting Ma,[7] Andrey Bobkov,[7] Eduard Sergienko,[7] Michael R. Jackson,[7] Branko Stefanovic,[8] Tatiana Kisseleva,[2,5] Gene W. Yeo,[3,4,5,9,10]* and David A. Brenner[1,7,12]*

[1]Department of Medicine, University of California San Diego, La Jolla, CA, 92093, USA
[2]Department of Surgery, University of California San Diego, La Jolla, CA, 92093, USA
[3]Department of Cellular and Molecular Medicine, University of California San Diego, La Jolla, California 92037, USA
[4]Institute for Genomic Medicine, University of California San Diego, La Jolla, CA, 92037, USA
[5]Sanford Stem Cell Institute, University of California San Diego, La Jolla, CA, 92037, USA
[6]Center for Computational Biology and Bioinformatics, University of California San Diego, La Jolla, CA, 92093, USA
[7]Sanford Burnham Prebys Medical Discovery Institute, La Jolla, CA 92037, USA
[8]Department of Biomedical Sciences, College of Medicine, Florida State University, Tallahassee, FL, USA
[9]Sanford Laboratories for Innovative Medicine, La Jolla, CA 92037, USA
[10]Center for RNA Technologies and Therapeutics, University of California San Diego, La Jolla, CA, 92037, USA
[11]These authors contributed equally
[12]Lead contact

Figures S1-S9 and Table S3

### Supplementary figures

A

| Patient ID | Pathology | Age | BMI | Gender | Alcohol (>2 drinks/day) | Steatosis grade, % | Lobular inflammation | Ballooning | Portal inflammation | Fibrosis stage | MASH/CRN score |
|---|---|---|---|---|---|---|---|---|---|---|---|
| D1 | NORMAL | 56 | 24 | M | No | 0 (<5%) | 0 | 0 | 0 | 0 | 0 |
| D2 | NORMAL | 6 | 25 | M | No | 0 | 0 | 0 | 0 | 0 | 0 |
| D3 | NORMAL | 34 | 22 | F | No | 0 | 0 | 0 | 1 | 0 | 0 |
| D4 | NORMAL | 42 | 32 | M | No | 0 | 0 | 0 | 1 | 0 | 0 |
| D5 | NORMAL | 42 | 27 | M | No | 0 | 0 | 0 | 0 | 0 | 0 |
| D6 | MASL | 70 | 38 | F | No | 0 (<5%) | 1 | 0 | 1 | 0 | 1 |
| D7 | MASL | 42 | 53 | F | No | 2 (50%) | 1 | 0 | 1 | 0 | 3 |
| D8 | MASL | 65 | 25 | M | No | 0 (<5%) | 2 | 0 | 1 | 0 | 2 |
| D9 | MASL | 62 | 39 | M | No | 1 (20%) | 1 | 0 | 1 | 0 | 2 |
| D10 | MASH | 55 | 44 | F | No | 1 (6-10%) | 1 | 1 | 2 | 4 | 3 |
| D11 | MASH | 50 | 30 | F | No | 3 (70%) | 2 | 0 | 1 | 2 | 5 |
| D12 | MASH | 54 | 29 | F | No | 0 (<5%) | 1 | 0 | 2 | 3 | 1 |
| D13 | MetALD | 47 | 31 | F | Yes | 3 (95%) | 1 | 1 | 1 | 4 | 5 |
| D14 | MetALD | 37 | 32 | M | Yes | 3 (95%) | 1 | 1 | 1 | 4 | 5 |
| D15 | MetALD | 36 | 26 | M | Yes | 3 (80%) | 1 | 2 | 1 | 3 | 6 |
| D16 | MetALD | 56 | 35 | M | Yes | 2 (50%) | 2 | 2 | 2 | 3 | 6 |
| D17 | MetALD | 44 | 25 | F | Yes | 0 (<1%) | 1 | 0 | 1 | 4 | 1 |
| D18 | MetALD | 52 | 32 | M | Yes | 2 (50%) | 1 | 0 | 2 | 3 | 3 |

B

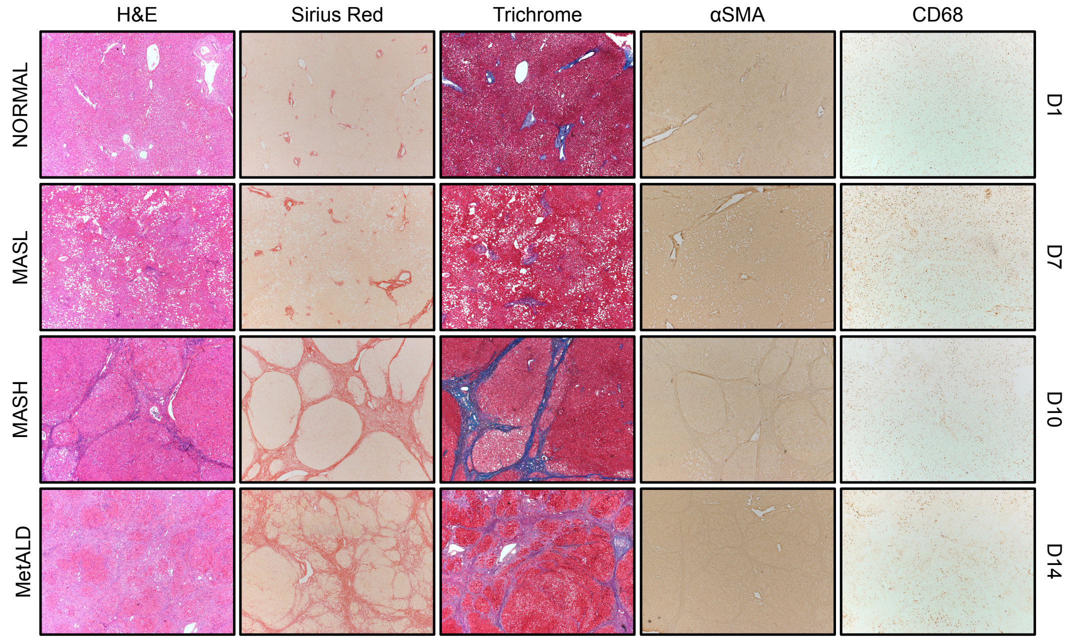


C

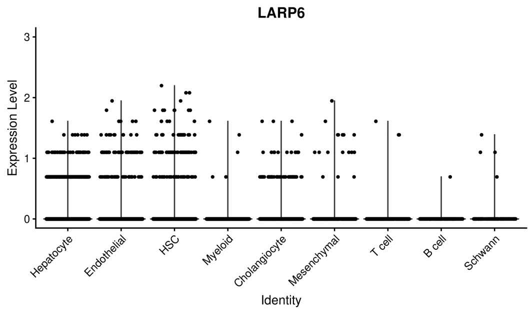


D

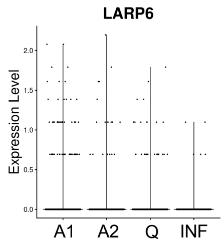


E

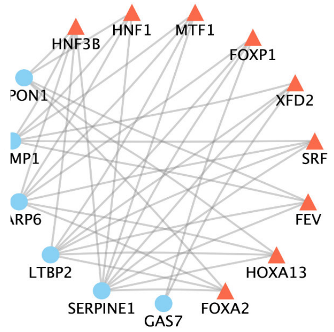

**Figure S1. LARP6 is expressed in activated HSCs of the fibrotic livers**

**(A)** Characterization of human livers selected for snRNA-seq and snATAC-seq. 4 distinct liver diagnoses of NORMAL, MASL, MASH, and MetALD were defined by a combination of pathological examination and alcohol consumption history. **(B)** Representative images of the livers stained with H&E, Sirius-Red, Masson's trichrome, anti-αSMA, anti-CD68 antibodies (×4 objective). **(C)** Violin plot of the LARP6 expression in the integrated dataset of liver cells from all donors. **(D)** Violin plot of the LARP6 expression in the integrated dataset of HSCs from all donors. **(E)** Network of transcription factors which are known to target LARP6.

A

| Patient ID | Pathology | Age | BMI | Gender | Alcohol (>2 drinks/day) | Steatosis grade, % | Lobular inflammation | Ballooning | Portal inflammation | Fibrosis stage | MASH/CRN score |
|---|---|---|---|---|---|---|---|---|---|---|---|
| D19 | MASH | 53 | 48 | F | No | 2 (34-66%) | 1 | 0 | 1 | 1 | 3 |
| D20 | MetALD | 37 | 28 | M | Yes | 3 (80%) | 1 | 2 | 0 | 3 | 6 |
| D21 | MetALD | 36 | 26 | M | Yes | 3 (80%) | 1 | 2 | 1 | 3 | 6 |
| D22 | MASH | 69 | 28 | F | No | 1 (20%) | 1 | 2 | 1 | 2 | 4 |
| D23 | NORMAL | 65 | 34 | M | No | 0 | 0 | 0 | 2 | 0 | 0 |

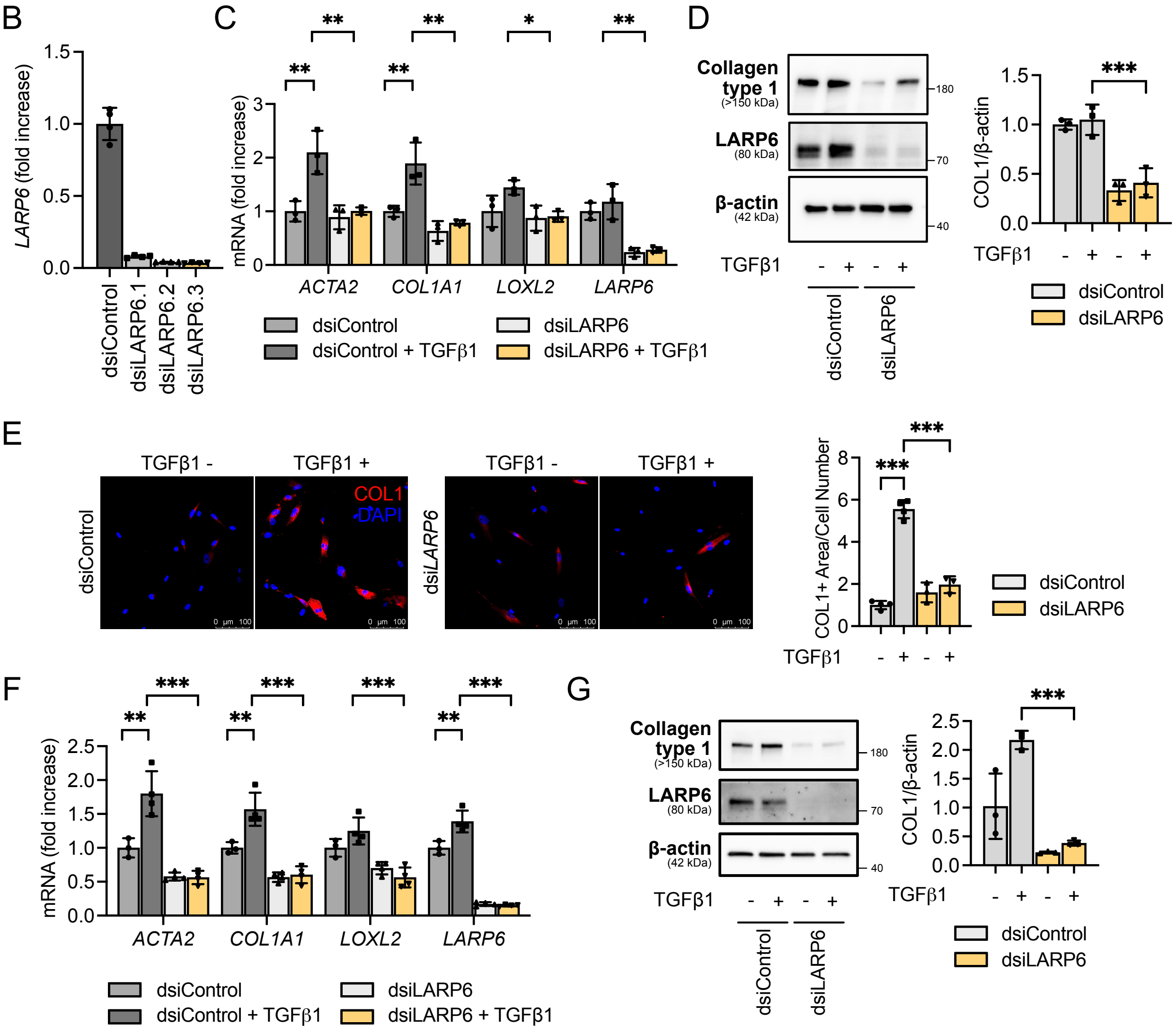


**Figure S2. Knockdown of LARP6 inhibits fibrogenic markers in human HSCs**
**(A)** Characterization of human donor livers used for cell isolation. HSCs were isolated from the fibrotic livers of donors D19-D21. Non-parenchymal cells (NPCs) were isolated from the liver of donor D22, and hepatocytes were isolated from the liver of donor D23. **(B)** Knockdown efficiency of human HSCs (donor D19) treated with three different LARP6-targeting dsiRNA.

dsiLARP6.3 was selected for further experiment. **(C-E)** Cultured HSCs (donor D20) were transfected for 48 h with dsiRNA and stimulated with TGFβ1 (5 ng/ml) for 24 h. The expression of fibrogenic genes was measured in dsiLARP6 (vs. dsiControl)-transfected HSCs ± TGFβ1 at the **(C)** mRNA and **(D)** protein levels. **(E)** dsiRNA-transfected HSCs ± TGFβ1 were stained with an anti-collagen type 1 (COL1) antibody, and the COL1-positive area was calculated and normalized by the cell number counted using DAPI. **(F, G)** Cultured HSCs (donor D21) were transfected for 48 h with dsiRNA and stimulated with TGFβ1 (5 ng/ml) for 24 h. The expression of fibrogenic genes was measured in LARP6-targeting dsiRNA (vs. dsi-negative control)-transfected HSCs ± TGFβ1 at the **(F)** mRNA and **(G)** protein levels. Data are mean ± SD; $^*P < 0.05$, $^{**}P < 0.01$, and $^{***}P < 0.001$, one-way ANOVA followed by Tukey's test.

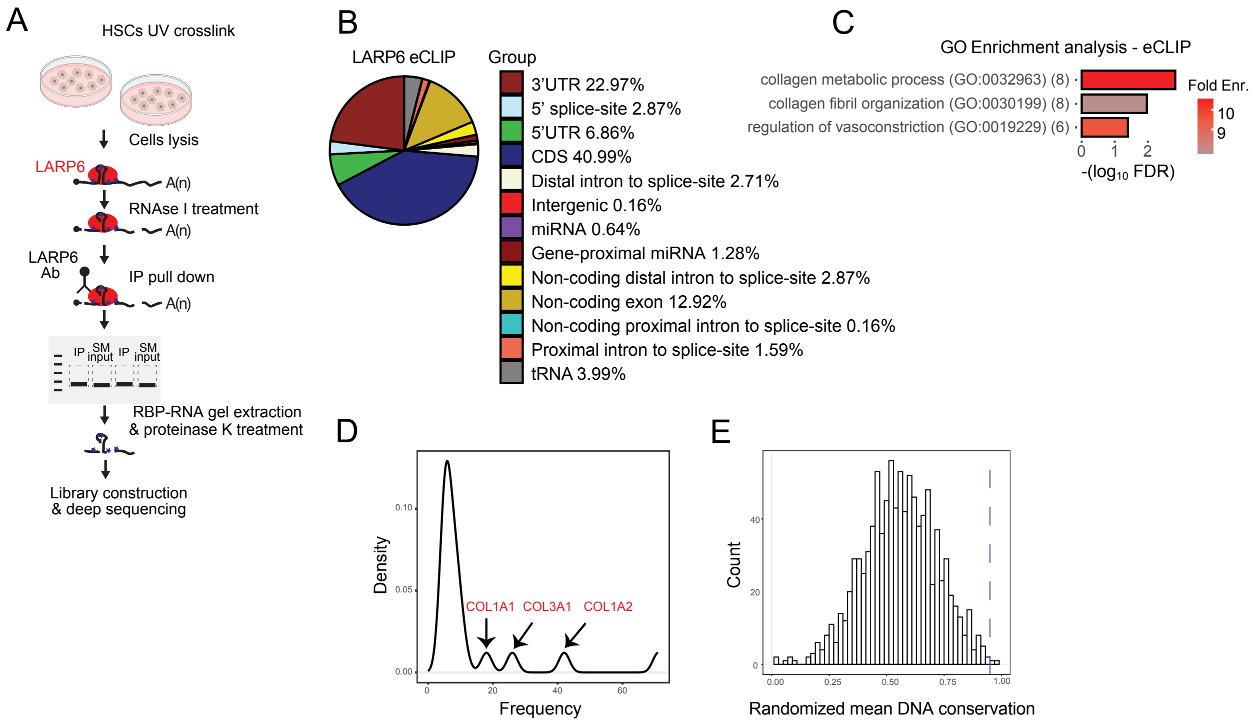


**Figure S3. LARP6 interacts with mature collagen mRNAs.**

**(A)** Schematic illustration of eCLIP on LARP6 in HSCs. **(B)** LARP6 eCLIP binding distribution in HSCs. **(C)** GO enrichment analysis using significant eCLIP peaks. RNA-seq genes with minimum of 30 reads were used as background for enrichment analysis. **(D)** Density of eCLIP analysis of LARP6 targets with minimum of 5 peaks. Peaks in *COL1A1*, *COL3A1* and *COL1A2* are demonstrated with black arrows. **(E)** DNA conservation analysis of 30 bp in 5' UTR and 30 bp in coding sequence was calculated using phyloP100way. The average for *COL1A1*, *COL1A2* and *COL3A1* is marked with a vertical dashed line, and the distribution of a random group composed of three genes was calculated a thousand times and represented with random distribution analysis. Collagen genes Z score = 15.39.

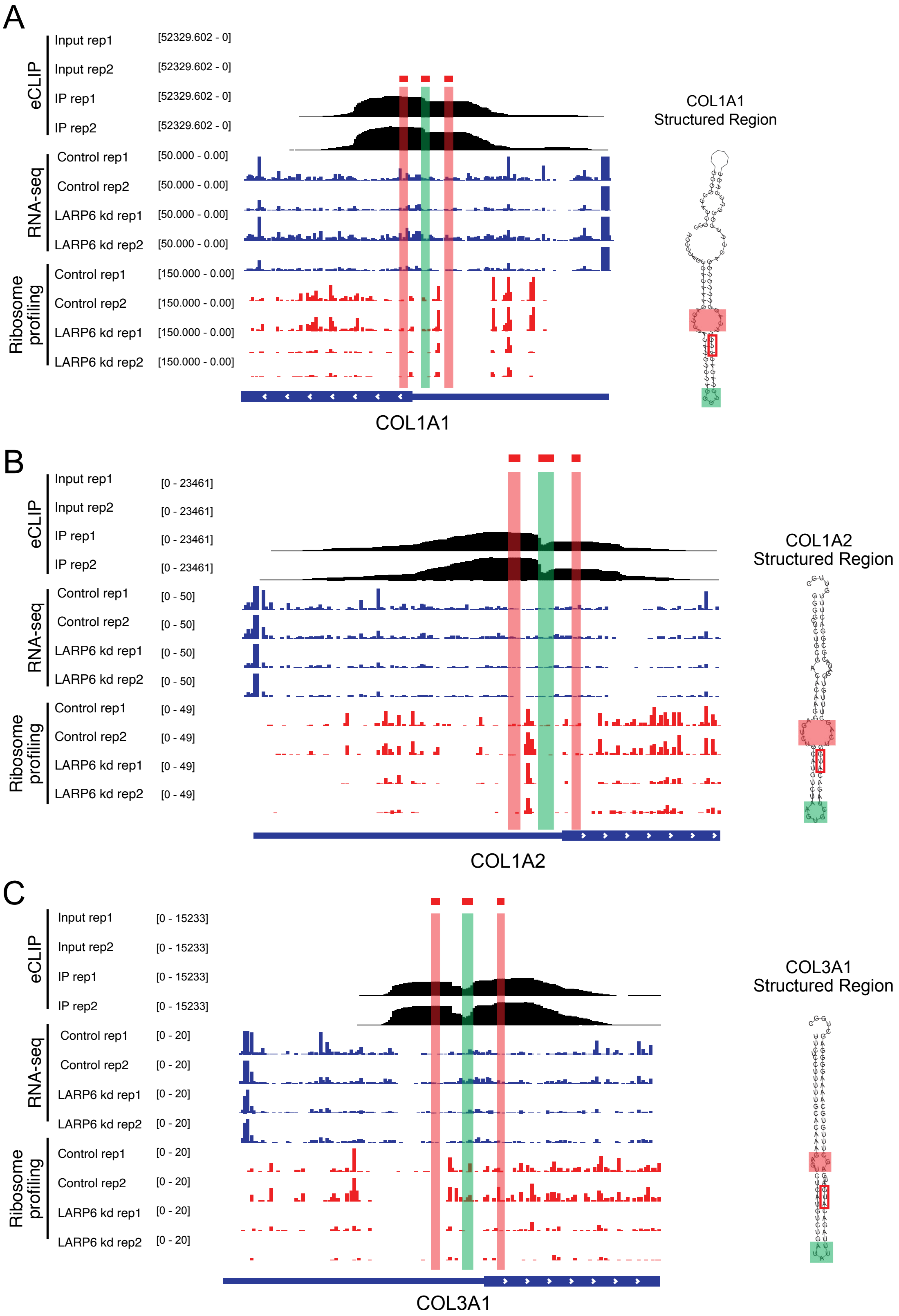
A
eCLIP
Input rep1 [52329.602 - 0]
Input rep2 [52329.602 - 0]
IP rep1 [52329.602 - 0]
IP rep2 [52329.602 - 0]
RNA-seq
Control rep1 [50.000 - 0.00]
Control rep2 [50.000 - 0.00]
LARP6 kd rep1 [50.000 - 0.00]
LARP6 kd rep2 [50.000 - 0.00]
Ribosome profiling
Control rep1 [150.000 - 0.00]
Control rep2 [150.000 - 0.00]
LARP6 kd rep1 [150.000 - 0.00]
LARP6 kd rep2 [150.000 - 0.00]
COL1A1
COL1A1 Structured Region
B
eCLIP
Input rep1 [0 - 23461]
Input rep2 [0 - 23461]
IP rep1 [0 - 23461]
IP rep2 [0 - 23461]
RNA-seq
Control rep1 [0 - 50]
Control rep2 [0 - 50]
LARP6 kd rep1 [0 - 50]
LARP6 kd rep2 [0 - 50]
Ribosome profiling
Control rep1 [0 - 49]
Control rep2 [0 - 49]
LARP6 kd rep1 [0 - 49]
LARP6 kd rep2 [0 - 49]
COL1A2
COL1A2 Structured Region
C
eCLIP
Input rep1 [0 - 15233]
Input rep2 [0 - 15233]
IP rep1 [0 - 15233]
IP rep2 [0 - 15233]
RNA-seq
Control rep1 [0 - 20]
Control rep2 [0 - 20]
LARP6 kd rep1 [0 - 20]
LARP6 kd rep2 [0 - 20]
Ribosome profiling
Control rep1 [0 - 20]
Control rep2 [0 - 20]
LARP6 kd rep1 [0 - 20]
LARP6 kd rep2 [0 - 20]
COL3A1
COL3A1 Structured Region

**Figure S4. LARP6 binds the structural elements of collagen mRNAs at the stem-bulge region.**

eCLIP, RNA-seq, and ribosome profiling of **(A)** *COL1A1*, **(B)** *COL1A2*, and **(C)** *COL3A1* are represented by black, blue, and red bars, respectively. eCLIP of TGFβ1-stimulated HSCs, RNA-seq and ribosome profiling of control and LARP6 knockdown (kd) HSCs, are shown. The stem bulge is marked with a red window, and the hairpin is indicated by a green window in the genomic (left) and structural modeling (right) of the collagen genes. The canonical AUG is marked with a red rectangle.

A

Consensus structural element of collagen RNAs

```
____CCCUCAGACACAAAGAGUCU_CAUGUCUAAGAUCUAGACAUGAUCAGCUUUGUGGAAAAGCGGACCUUGCU_C________________
......((((((((((((.....((((((((((...))))))))))....))))))))...))))).......................... (-24.84 = -24.07 +  -0.77) [sci = 0.7535]

        ......((((((((((((.....((((((((((...))))))))))....)))))))).
col1a1   ----TCCCCAGCCACAAAGAGTCTACATGTCTAGGGTCTAGACATGTTCAGCTTTGTGGA 56
col1a2   GGGGCTCTGCGACACAAGGAGTCTGCATGTCTAAGTGCTAGACATGCTCAGCTTTGTGGA 60
col3a1   TTCTCCTTTTTGCACAAAGAGTCT-CATGTCTGATATTTAGACATGATGAGCTTTGTGCA 59
         .........10........20........30........40........50........6

         ..))))).............................
col1a1   CCTCCGGCTCCTGCTCCTCTTAGCGGCCACCGCC 90
col1a2   TACGCGGACTTTGTTGC----------------- 77
col3a1   AAAGGGGAGCTGGC-------------------- 73
         0........70........80........90...
```

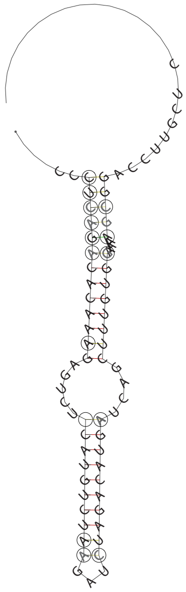

B

COL1A1

```
consensus sequence    UCCCCAGCCACAAAGAGUCUACAUGUCUAAGAUUUAGACAUGUUCAGCUUUGUGGAUAUCCGGCUCCUGCUCCUCAUAGCAGCCACUGCC
                      .......((((((((.....((((((((((...))))))))))....))))))))...........(((((......))))).........
                      MEA(-23.57 = -23.90 +   0.33) [sci = 0.7915]

Homo sapiens          UCCCCAGCCACAAAGAGUCUACAUGUCUAGGGUCUAGACAUGUUCAGCUUUGUGGACCUCCGGCUCCUGCUCCUCUUAGCGGCCACCGCC
Mus musculus          ACCCCAGCCGCAAAGAGUCUACAUGUCUAGGGUCUAGACAUGUUCAGCUUUGUGGACCUCCGGCUCCUGCUCCUCUUAGGGGCCACUGCC
Gallus gallus         CAAAGAGCCCCAAAGAGUCUACAUGUCUAAUAUUUAGACAUGUUCAGCUUUGUGGAUUCUCGGUUACUGCUGUUGAUAGCAGCGACUGUA
Xenopus tropicalis    -AAAGGACCACAGAGAGUCUACAUGUCUAU---UUAGACAUGUUCAGCUUUGUGGAUACACGGACCCUGCUACUUAUAGCAGCAACAAUC
Danio rerio           UCCCUCGCCUCAAAGAGUCUACAUGUCUAAUAUUUAGACAUGUUCAGCUUUGUGGAUAUUCG---------------------------
                       ...  .** **.***************     *********************    **
```

C

COL1A2

```
consensus sequence  GGGGCUCUGCAACACAAGGAGUCUGCAUGUCUAACUGCUAGACAUGCUCAGCUUUGUGGAUACGCGGACUCUGUUGCUGCUUGC
                    ............(((((((.....(((((((((.....)))))))))....))))))).....((((.(......))))).....
                    MEA(-23.46 = -23.66 +   0.20) [sci = 0.7985]

Homo sapiens        GGGGCUCUGCGACACAAGGAGUCUGCAUGUCUAAGUGCUAGACAUGCUCAGCUUUGUGGAUACGCGGACUUUGUUGCUGCUUGC
Mus musculus        GGGGCUCUGCAACACAAGGAGUCUGCAUGUCUAAGUGGUAGACAUGCUCAGCUUUGUGGAUACGCGGACUCUGUUGCUGCUUGC
Gallus gallus       GGGGCUCUGCAACACAAGGAGUCUGCAUGUCUAGCAAGUAGACAUGCUCAGCUUUGUGGAUACGCGGAUUUUGUUGCUGCUCGC
Xenopus tropicalis  GG--CUCUCACUCACAAGGAGUCUGCAUGACUAGCCGCUAGCCAUGCUCAGCUUUGUGGAUAUGCGGACUGUGCUGCUGCUGGC
Danio rerio         AGGGCACUGCGGCACAAGGAGUCUGCAUGUCGG---UUUAGACAUGCUCAGCUUUGUGGAUACCCGGAUUCUGUUGCUGCUUGC
                    .*  *:** .  *****************:* .     ***.*******************  **** * ** ******* **
```

D

COL3A1

```
consensus sequence  UUUCCUUUCCCCCCUUUU________GCACAAAGAGUCUCAUGUCUAAUAUUUAGACAUGAUGAGCUUUGUGCAAAAUGGAACCUGGC
                    ..........................((((((((..(((((((((((((...)))))))))))).)).)))))))).............
                    MEA(-25.27 = -24.77 +  -0.50) [sci = 0.8030]

Homo sapiens        UU--CUUUUCUCCUUUUU--------GCACAAAGAGUCUCAUGUCUGAUAUUUAGACAUGAUGAGCUUUGUGCAAAAGGGGAGCUGGC
Mus musculus        CUUCCUCUCUCCUCUUUU--------GCACAAAGAGUCUCAUGUCUGAUAUUUAGACAUGAUGAGCUUUGUGCAAAGUGGAACCUGGU
Gallus gallus       UUUCUUUCCCCCCCAUUUUUCUUUUUGCACAGAGAGUCUCAUGUCUAAUAUUUAGACAUGAUGAGCUUUGUGCAAAAAGUAAGUUUGU
Xenopus tropicalis  UAUUUUGUUUUCCCCCCG---AGUCUGCACUUAGAGUCUCAUGUCUAACAUUUAGACAUGAUGAGCUUAGUGCAAAUUGCGUCCCUUC
                     :   *     *              ****: *************.* ******************:*******  * .:
```

**Figure S5. Structural elements of collagen mRNAs are highly conserved in mammals.** **(A)** MEA (maximum expected accuracy) structure sequence and structural conservation index (SCI) of *COL1A1*, *COL1A2* and *COL3A1* was calculated with RNAalifold using ViennaRNA package. **(B-D)** MEA structure and SCI was calculated with RNAalifold using ViennaRNA package for **(B)** *COL1A1*, **(C)** *COL1A2* and **(D)** *COL3A1*. Canonical AUG is marked with black rectangle. *COL3A1* is not expressed in Danio rerio genome and not represented[66].

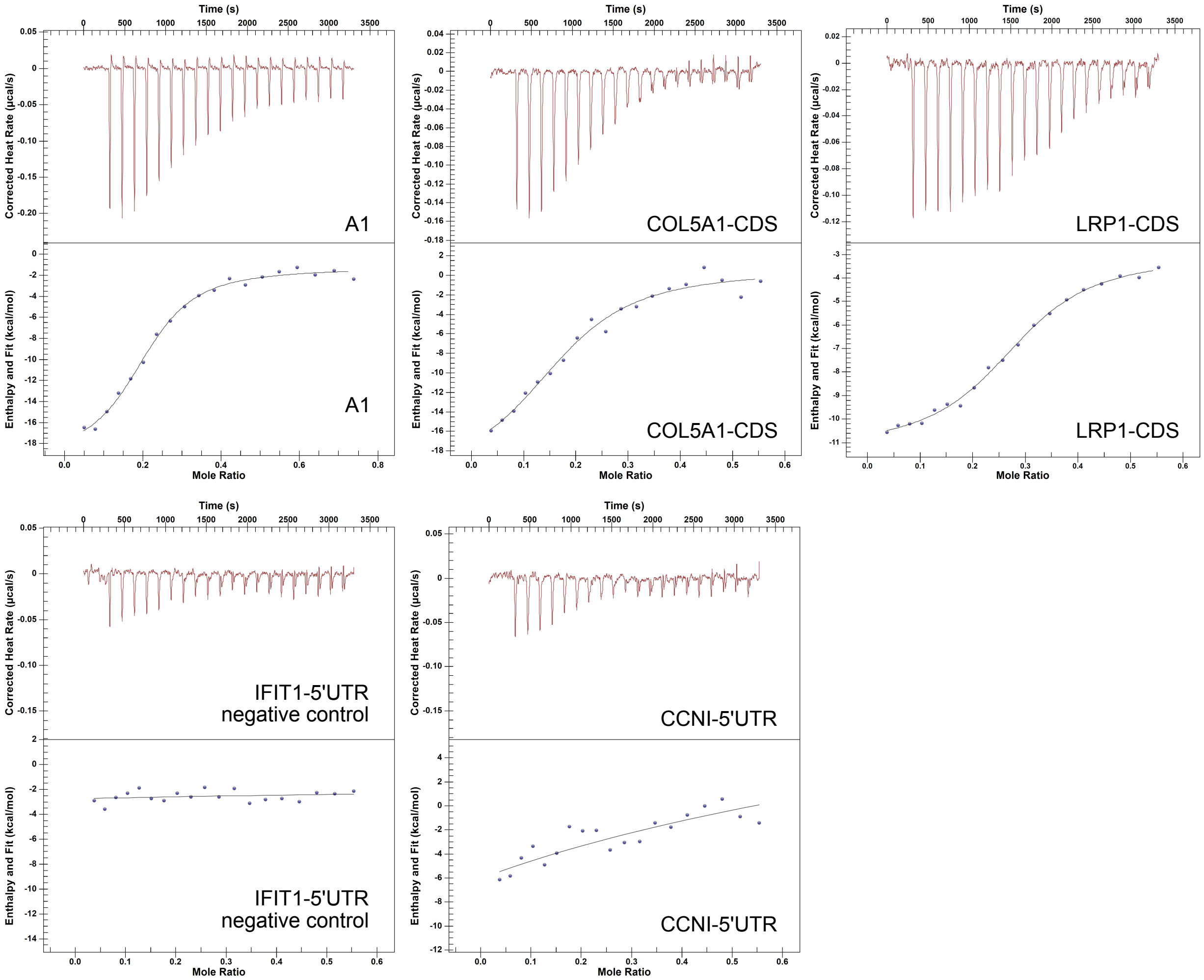


**Figure S6. ITC curves of LARP6 binding to different target RNAs.**
ITC curves of LARP6 binding to A1, *COL5A1*-CDS, *LRP1*-CDS, *CCNI*-5'UTR, and IFIT1-5'UTR.

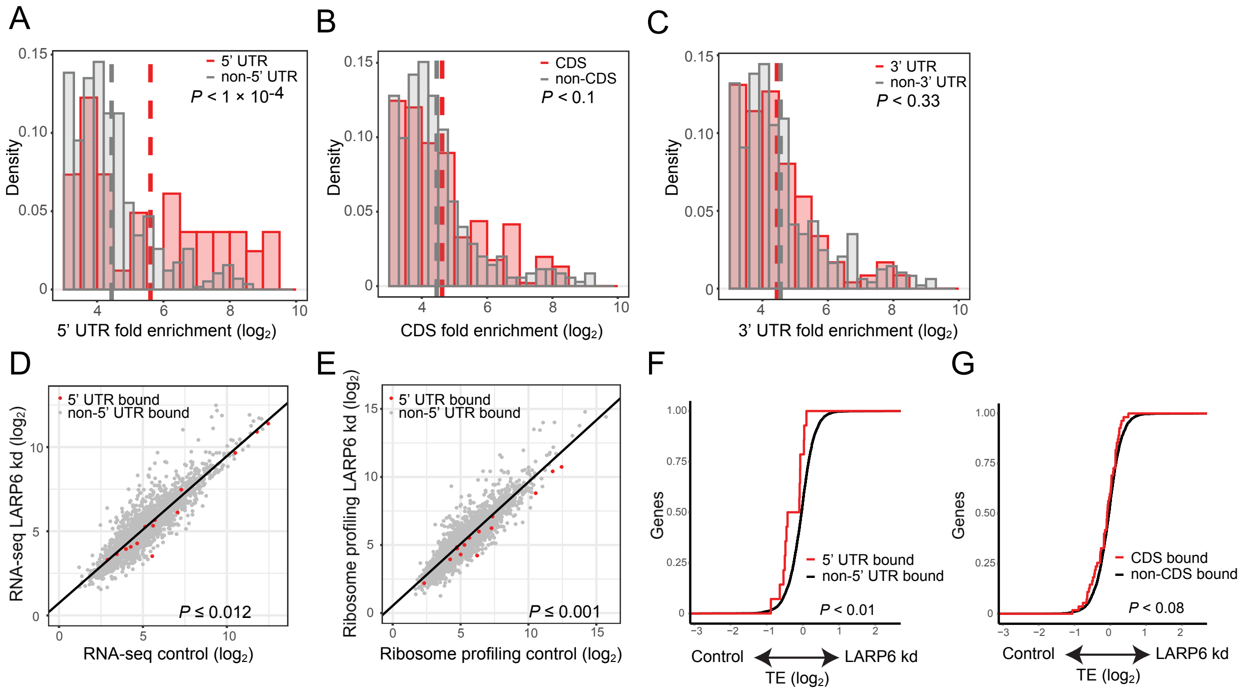


**Figure S7. LARP6 directly regulates translation via binding to 5'UTRs.**
**(A-C)** Density analysis of fold enrichment ($\log_2$) from HSC eCLIP analysis. **(A)** Non-5' UTR peaks and 5' UTR peaks are represented with grey and red bars, respectively, and mean is indicated with a dashed line. Same analysis for **(B)** CDS and **(C)** 3' UTR targets. *P* value was calculated using student *t*-test. **(D-E)** Correlation of **(D)** RNA-seq and **(E)** ribosome profiling data in control and LARP6 kd HSCs. 5' UTR targets of LARP6 from HSCs eCLIP data are marked in red dots while the non-bound targets are marked with grey dots. *P* value is represented from student *t*-test. **(F-G)** Cumulative translation efficiency was calculated with the ratio of RNA-seq and ribosome profiling data in LARP6 kd cells to control. The **(F)** 5' UTR and **(G)** CDS targets from eCLIP data are marked with red line, and the non-bound targets with a black line. *P* value was calculated with a student *t*-test.

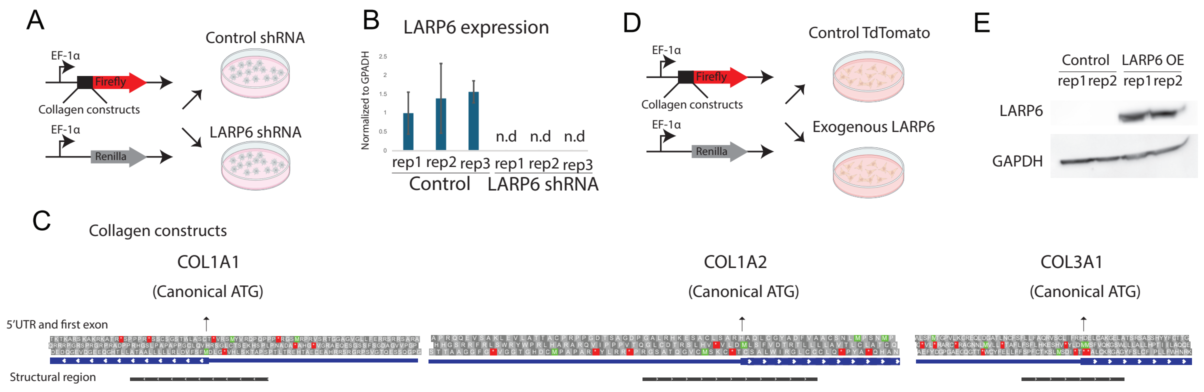


**Figure S8. LARP6 is necessary and sufficient for translation regulation.**
**(A)** Schematic illustration of LARP6 knockdown (kd) using shRNA in HSCs. **(B)** qPCR for LARP6 expression in wild-type (control) or LARP6 knockdown (LARP6 shRNA) human HSCs (donor D22), normalized to GAPDH expression. n.d., not detected. **(C)** Schematic illustration of collagen 5' UTR and first coding exon. Structural elements are represented with black bars. **(D)** Schematic illustration of exogenous LARP6 in HeLa cells. **(E)** Western blot analysis of LARP6 and GAPDH expression in either exogenous tdTomato or LARP6 in HeLa cells.

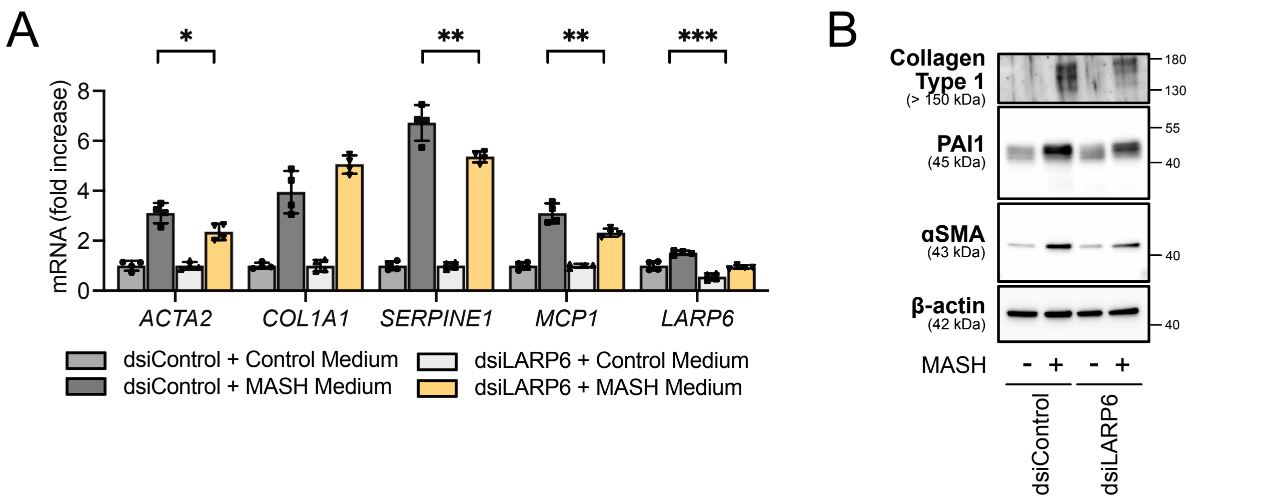


**Figure S9. HSC-specific knockdown of LARP6 inhibits MASH-induced fibrosis in human liver spheroids.**

Human liver spheroids were generated using hepatocytes (donor D23), NPCs (donor D22), and dsiLARP6-transfected HSCs (donor D19) and incubated under MASH conditions. **(A)** Fibrogenic markers in liver spheroids was assessed using qPCR. **(B)** Western blotting was performed to measure Collagen Type 1, PAI-1, and αSMA expression.

**Supplemental table**

**Table S3.** Oligonucleotides used in this study, related to STAR Methods.

| Primer for qPCR | | |
|---|---|---|
| Gene | Sequence 5'→3' | |
| **Human** | **Forward primer** | **Reverse primer** |
| *SERPINE1* | AGTGGACTTTTCAGAGGTGGA | GCCGTTGAAGTAGAGGGCATT |
| *HPRT* | CCTGGCGTCGTGATTAGTGAT | AGACGTTCAGTCCTGTCCATAA |
| *ACTA2* | CACCATCGGAAATGAACGTTT | GACTCCATCCCGATGAAGGA |
| *COL1A1* | AAGAGGAAGGCCAAGTCGAG | CACACGTCTCGGTCATGGTA |
| *LOXL2* | AGGACATTCGGATTCGAGCC | CTTCCTCCGTGAGGCAAAC |
| *COL1A2* | CCGTGCTTCTCAGAACATCA | CTTGCCCCATTCATTTGTCT |
| *TIMP1* | CTTCTGCAATTCCGACCTCGT | ACGCTGGTATAAGGTGGTCTG |
| *LARP6* | GGAGGTGAGAATGAGCGTGAG | GTGTTTTAGCAAAAAGGCGTCC |
| *LARP6* | TGAAAACCTGGAGAAGGACGCC | ATGTGCTGTGGTTCTCCAGTCC |
| *18srRNA* | GTAACCCGTTGAACCCCATT | CCATCCAATCGGTAGTAGCG |
| *GAPDH* | CCACCCATGGCAAATTCC | TGGGATTTCCATTGATGACAAG |

| RNA oligos for FRET and ITC analysis | | | | | |
|---|---|---|---|---|---|
| Gene | Sequence | Chr | Start | End | Strand |
| *CCNI* | GGGUCAUCAUGGAACUAAUUC GCUGACCGACCCAGCGGCCGC AGCCGUGCGUCCCGCUCG | chr 4 | 77075667 | 77075727 | - |
| *LRP1* | CUGCAUGUGCACAGCCGGCUA UAGCCUCCGGAGUGGCCAGCA GGCCUGCGAGGGCGUAGG | chr 12 | 57181238 | 57181292 | + |
| *COL5A1* | CUGCUGCUGCUGCUGUGGGCG CCGCCUCCGAGCCGCGCAGCU CAGCCAGCAGAUCUCCUG | chr 9 | 134642256 | 134690931 | + |
| *COL1A1* | CCAGCCACAAAGAGUCUACAUG UCUAGGGUCUAGACAUGUUCA GCUUUGUGGACCUCCGG | chr 17 | 50201489 | 50201549 | - |

| | | | | | |
|---|---|---|---|---|---|
| *IFIT1* | AGACAGAAUAGCCAGAUCUCAGAGGAGCCUGGCUAAGCAAAACCCUGCAGAACGGCUGCC | chr 10 | 89392637 | 89392697 | + |
| A1 | CCACAAAGAGUCUACAUGUCUAGGGUCUAGACAUGUUCAGCUUUGUGG | chr 17 | 50201497 | 50201545 | - |

| Sequences for reporter elements | |
|---|---|
| Gene | Sequence |
| 5’UTR targets | |
| *CCDC85B* | CACTGCCTGCCGCGTGCGGAGCCGGAGCCCGAGCCTGAGTGGCGCCGGGCCCGACGTGGGGCTCCTGGGCCGCGGCGGCGGGCGGGCGATGCTCCAGAGGCCTGACCAGCC |
| *COL1A1* | GCAGACGGGAGTTTCTCCTCGGGGTCGGAGCAGGAGGCACGCGGAGTGTGAGGCCACGCATGAGCGGACGCTAACCCCCTCCCCAGCCACAAAGAGTCTACATGTCTAGGGTCTAGAC |
| *COL1A2* | AGCACCACGGCAGCAGGAGGTTTCGGCTAAGTTGGAGGTACTGGCCACGACTGCATGCCCGCGCCCGCCAGGTGATACCTCCGCCGGTGACCCAGGGGCTCTGCGACACAAGGAGTCTGCATGTCTAAGTGCTAGAC |
| *COL3A1* | GGCTGAGTTTTATGACGGGCCCGGTGCTGAAGGGCAGGGAACAACTTGATGGTGCTACTTTGAACTGCTTTTCTTTTCTCCTTTTTGCACAAAGAGTCTCATGTCTGATATTTAGAC |
| *MAP4K4* | ACTCGCTCAACTCGGCGCCGCCGCGGCCCCACGCTCCGGGCCCGTCCTCGAGGCGCGCGGCGCGGGGCGCGGGCGCCGGGGCCTGAGGCGGCGGGCGACGCCCGGGGGCCTGACGGCCGGCCCCGCGCCATGGTGTGAGCGCCGCCGCCCGTGCACGCTCCGTCCGCCCTCCGCGCGGCCCGGCCGGCAGAGAGCCCCGAGCGGCCCGAGAGCGCAGCCGAGCCCGCCGCCGCCGCCCGCGGCCCCGCGAGGAGAGTACCGGGCCGGCTCGGCTGCCGCGCGAGGAGCGCGGTCGGCGGCCTGGTCTGCGGCTGAGATACACAGAGCGACAGAGACATTTATTGTTATTTGTTTTTTGGTGGCAAAAAGGGAAA |
| *SPTBN1* | AGTCCCTCCCTCGGCCGCCTCTCCTCCCGGAGCGAGCGCGCAGCCCTGCGCAGCAGCGCCCACTGGTCCCGTCCTGTGAGCCCCGGCCCCAGCCGCGGACAGACCCGCGGAGTCGCCTCCCGGCCCACCCGCCCGGCCGCCGAGGAGCGGGAGGAGGACGGGACCCCGGCGCCCCCACCCCATCCCCGGGAGAACTCTAAGAAGGAGCTGATGTGGAGGAGCAGCTGAGACAGTTCAAG |

| Collagen targets | | |
|---|---|---|
| Element | Wild type | Remove canonical ATG |
| COL1A1 structural element | TCCCCAGCCACAAAGAGTCTACATGTCTAGGGTCTAGACATGTTCAGCTTTGTGGACCTCCGGCTCCT | TCCCCAGCCACAAAGAGTCTACATGTCTAGGGTCTAGACTTCAGCTTTGTGGACCTCCGGCTCCTGCTCCTCTTA |

| | | |
|---|---|---|
| | GCTCCTCTTAGCGGCCACCGCC | GCGGCCACCGCC |
| COL1A2 structural element | GGGGCTCTGCGACACAAGGAGTCTGCATGTCTAAGTGCTAGACATGCTCAGCTTTGTGGATACGCGGACTTTGTTGC | GGGGCTCTGCGACACAAGGAGTCTGCATGTCTAAGTGCTAGACCTCAGCTTTGTGGATACGCGGACTTTGTTGC |
| COL3A1 structural element | TTCTCCTTTTTGCACAAAGAGTCTCATGTCTGATATTTAGACATGATGAGCTTTGTGCAAAAGGGGAGCTGGC | TTCTCCTTTTTGCACAAAGAGTCTCATGTCTGATATTTAGACAGCTTTGTGCAAAAGGGGAGCTGGC |
| shRNA oligos | | |
| LARP6 shRNA | CCGGGCACATGCTTTGAAGTATTCACTCGAGTGAATACTTCAAAGCATGTGCTTTTTTG | |
